# An expanded LPS translocon mediates surface lipoprotein export in the Bacteroidota

**DOI:** 10.64898/2026.10.03.756453

**Authors:** Xiaolong Liu(刘晓龙), Luis Orenday Tapia, Owain J. Bryant, Joseph J.E. Caesar, Hans Elmlund, Faraz K. Mardakheh, Susan M. Lea, Ben C. Berks

## Abstract

Cell surface lipoproteins (SLPs) play crucial roles in the pathogenesis and cell biology of gram-negative bacteria, yet the route by which they reach the cell surface has remained unknown except in special cases^1–3^. Here we show that in Bacteroidota bacteria, a phylum that includes prominent human commensals and anaerobic pathogens, SLP export is carried out by the outer membrane lipopolysaccharide (LPS) translocon. Isolation and structural characterisation of LPS translocon complexes from two Bacteroidota species reveals that they are massively enlarged and elaborated relative to the *Escherichia coli* paradigm with up to eight additional subunits located within, and on either side of, the outer membrane. We establish a role for the universally conserved core LptD subunit and four of the additional Bacteroidota-specific subunits in SLP export. Expansion of the LPS translocon in the Bacteroidota may allow export of the unusually complex SLPs found in this phylum. Our discoveries raise the possibility that SLP export is a conserved second function of the LPS transport pathway across gram-negative bacteria.

## Main text

The Outer Membrane (OM) of gram-negative bacteria is an asymmetric bilayer in which the outer leaflet is composed of LPS and the inner leaflet of phospholipids. OM-anchored lipoproteins are also asymmetrically distributed with distinct sets of lipoproteins associated with the periplasmic and extracellular sides of the membrane. Consequently, OM biogenesis requires mechanisms to transfer both LPS and cell surface lipoproteins into the extracellular leaflet of the membrane.

The mechanism by which LPS is trafficked to the OM has been well-characterized in *Escherichia coli*^4–6^. An ATP-driven transporter extracts LPS from the Inner Membrane (IM) and pushes it through a periplasmic bridge to a translocon in the OM where it is guided it into the extracellular leaflet^4,5^. The periplasmic bridge is proposed to be a continuous β-taco structure that extends from subunits of the IM transporter through the periplasmic protein LptA and into the translocon subunit LptD^7,8^. A deep hydrophobic groove running along the β-taco accommodates the acyl tails of the LPS molecules, shielding them from the aqueous periplasm and allowing LPS to slide along the bridge. In *E. coli* and other proteobacteria the LPS translocon has a conserved core formed by the proteins LptD and LptE^9–11^. LptD contains a kidney-shaped 26-strand transmembrane β-barrel with a weak seam between the first and last barrel strands that forms a lateral gate. The pore of the LptD barrel is plugged at the gate-distal end by the lipoprotein LptE and sealed at the gate-proximal end by extracellular loops from the barrel. The β-taco domain of LptD that forms the end of the periplasmic bridge is positioned outside the lateral gate. LPS molecules exiting the bridge are envisaged to trigger opening of both the lateral gate and the proximal end of the barrel pore allowing the polar oligosaccharide part of the LPS molecule to pivot through the pore to the extracellular side of the membrane while the acyl chains remain in the non-polar part of the membrane outside the gate^9,12,13^. In *E. coli* and some other Enterobacteriaceae the LPS translocon has recently been shown to include two additional, non-essential lipoproteins LptM^12,14–16^ and LptY/YedD^14,17^. SLPs are also initially assembled at the IM, where their N-terminal cysteine residue is tri-acylated. However, no widespread mechanism for their subsequent export has yet been identified^2^.

SLPs are particularly abundant and important in the Bacteroidota, a phylum that constitutes a major part of the human microbiome. Bacteroidota include commensals that play key roles in gut health as well as anaerobic pathogens responsible for sepsis and periodontitis. Bacteroidota characteristically possess hundreds of SLPs, a large proportion of which are involved in utilising extracellular polysaccharides. Individual polysaccharides are typically processed by multiple SLPs, including extracellular glycosyl hydrolases and components of SUS <u>s</u>ugar <u>u</u>ptake <u>s</u>ystems^18,19^. Diverse other uses of SLPs by Bacteroidota include as subunits of the β-barrel Assembly Machinery (BAM) used to insert transmembrane proteins into the OM^20,21^, for adhesion to hosts^22^ and as protective protein coats^23^.

The pathway by which SLPs reach the Bacteroidota cell surface is unknown. Various specialised SLP export systems identified in other phyla can mostly be excluded as candidates for this pathway. The Surface Lipoprotein Assembly Modulator (SLAM) is absent from Bacteroidota ^24^, the Type II secretion system is only sporadically present^25–27^, and both the BAM complex and the Translocation and Assembly Module (TAM) have previously been ruled out experimentally^20,21,28^. However, it has recently been reported that the spirochaete *Borrelia burgdorferi*, which lacks LPS, uses a highly divergent homologue of the LPS translocon protein LptD to export SLPs^29^ (though this conclusion has been challenged^30^). Although Bacteroidota possess a canonical LptD rather than one resembling the *B. burgdorferi* homologue, the precedent of SLP export by a LptD family protein raises the possibility that the Bacteroidota LPS translocon might also have the ability to transport SLPs. Consistent with this idea, the *E. coli* LPS translocon has been argued to adventitiously export some normally periplasmic lipoproteins^15^. A shared mechanism for LPS and SLP transport is mechanistically plausible because both substrates comprise a large hydrophilic moiety attached to multiple acyl chains.

### LptD is required for surface lipoprotein export in the Bacteroidota *F. johnsoniae*

To test the hypothesis that Bacteroidota SLP export is mediated by the LPS translocon we asked whether removing the core translocon component LptD (Fjoh_4586) from the free-living Bacteroidota *F. johnsoniae* interferes with SLP localisation to the cell surface. Because LptD is required for the essential process of LPS export, we constructed a strain in which LptD could be conditionally depleted by removal of the inducer anhydrotetracyline (aTC). Using this strain, we confirmed that LptD is essential for growth under standard laboratory conditions (Fig. 1a).

**Fig. 1:**
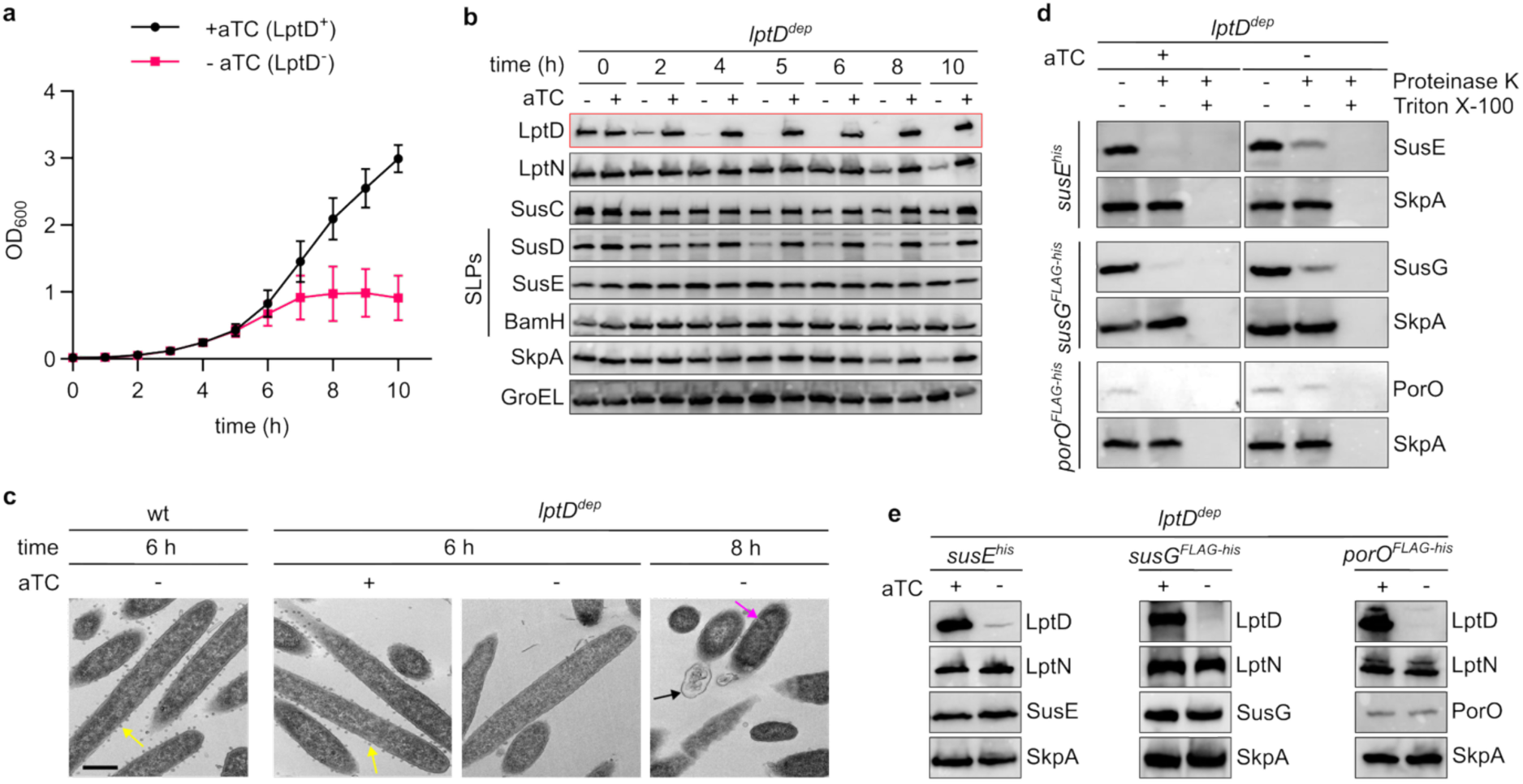
LptD is required for SLP export in *F. johnsoniae*. **a-c**, Phenotypic effects of LptD depletion. The LptD depletion strain (*lptD*^dep^) was cultured in rich (CYE) medium. In the -aTC samples the aTC inducer was removed at 0 h to initiate LptD depletion. wt, wild type. Similar data were obtained for three biological repeats. **a,** Growth curves (means ± 1 SD for four biological repeats). **b,** Whole cell immunoblots. SLPs, surface lipoproteins; SkpA, a periplasmic protein; GroEL a cytoplasmic protein. **c,** Representative transmission electron microscopy images during LptD depletion. Yellow arrows highlight budding OM vesicles, black arrows highlight OM blebbing, magenta arrows indicate internal membranous structures. Scale bar, 500 nm. **d,** Depletion of LptD for 6 h leads to intracellular accumulation of the SLPs SusE, SusG, and PorO as assessed by Proteinase K accessibility. Triton X-100 permeabilises the OM. Reactions were analysed by immunoblotting. SkpA, is a periplasmic protein. Similar data were obtained for three or more biological repeats. **e,** Whole cell immunoblots of the cells used in **d** confirming no changes in the levels of the substrate proteins or loss of LptN or the periplasmic marker protein SkpA upon LptD depletion. Similar data were obtained for three or more biological repeats.

LptD was fully depleted by 6 h after removal of the inducer (Fig. 1b) at which point cell growth slowed (Fig. 1a). There was no loss of periplasmic contents at this point (Fig. 1b, SkpA lanes) showing that the OM remained intact. Consistent with this, cells at this stage rarely exhibited morphological defects, although the production of OM vesicles (OMVs) was almost completely supressed (Fig. 1c). At later time points OM blebbing became common, and many cells developed irregular internal membranous structures or vacuoles resembling the envelope defects observed following LptD depletion in *E. coli*^31,32^. We therefore chose the 6 h time point to assess the effects of LptD loss on SLP biogenesis before the onset of substantial defects in OM integrity or cell morphology.

Immunoblotting against the three subunits of the most abundant *F. johnsoniae* SUS system revealed a marked reduction in the levels of the SLP SusD (Fjoh_0404) subunit, but not the membrane protein SusC (Fjoh_0403) or the SLP SusE (Fjoh_0405) at the 6 h depletion point (Fig. 1b). As all three proteins are encoded by the same operon, the selective loss of SusD is unlikely to result from a transcriptional effect and instead implicates LptD in the biogenesis or stability of this SLP. Because the levels of SusE and another SLP, BamH, were maintained, SLP loss is not a general consequence of LptD depletion.

To determine whether LptD depletion impairs SLP transport to the cell surface we used a protease protection assay to assess the surface exposure of representative, structurally diverse, SLPs at the 6 h depletion point. In these experiments protease added outside the cells digests SLPs on the cell surface, while any SLPs that remain inside the cell are protected from digestion because the protease cannot cross the OM. The SLPs tested were SusE, the glycosyl hydrolase SusG (Fjoh_1210), and a β-helix protein PorO (Fjoh_4720). Because SLPs are intrinsically protease-resistant, each test protein was fused to a protease-sensitive epitope tag to allow implementation of the protection assay.

All three test proteins exhibited substantial protease protection after LptD depletion whereas the same SLPs were fully susceptible to protease digestion in LptD-replete cells (Fig. 1d,e). Thus, LptD depletion impedes the transport of diverse SLPs to the cell surface, confirming the hypothesis that the LPS translocon mediates SLP export. Complete protease protection is not expected following LptD depletion because a proportion of the SLP molecules present at the 6 h assay point will have reached the cell surface before LptD levels became limiting for transport.

### Isolation of Bacteroidota LptD complexes

We investigated whether the Bacteroidota LPS translocon is modified relative to the *E. coli* paradigm in order to accommodate SLP export. To do this we isolated the endogenous *F. johnsoniae* LPS translocon (LPT_Fj_,) using an affinity tag fused to the native LptD protein and determined its structure by cryo-EM. In addition to LptD, the purified complex contains nine protein subunits (Fig. 2a). Two are the canonical LPT components LptA and LptE. We designate the remaining proteins LptN, LptP, LptQ, LptR, LptS, and LptT, with two copies of LptS present in the complex. No components of the canonical IM LPS transporter were detected.

**Fig. 2:**
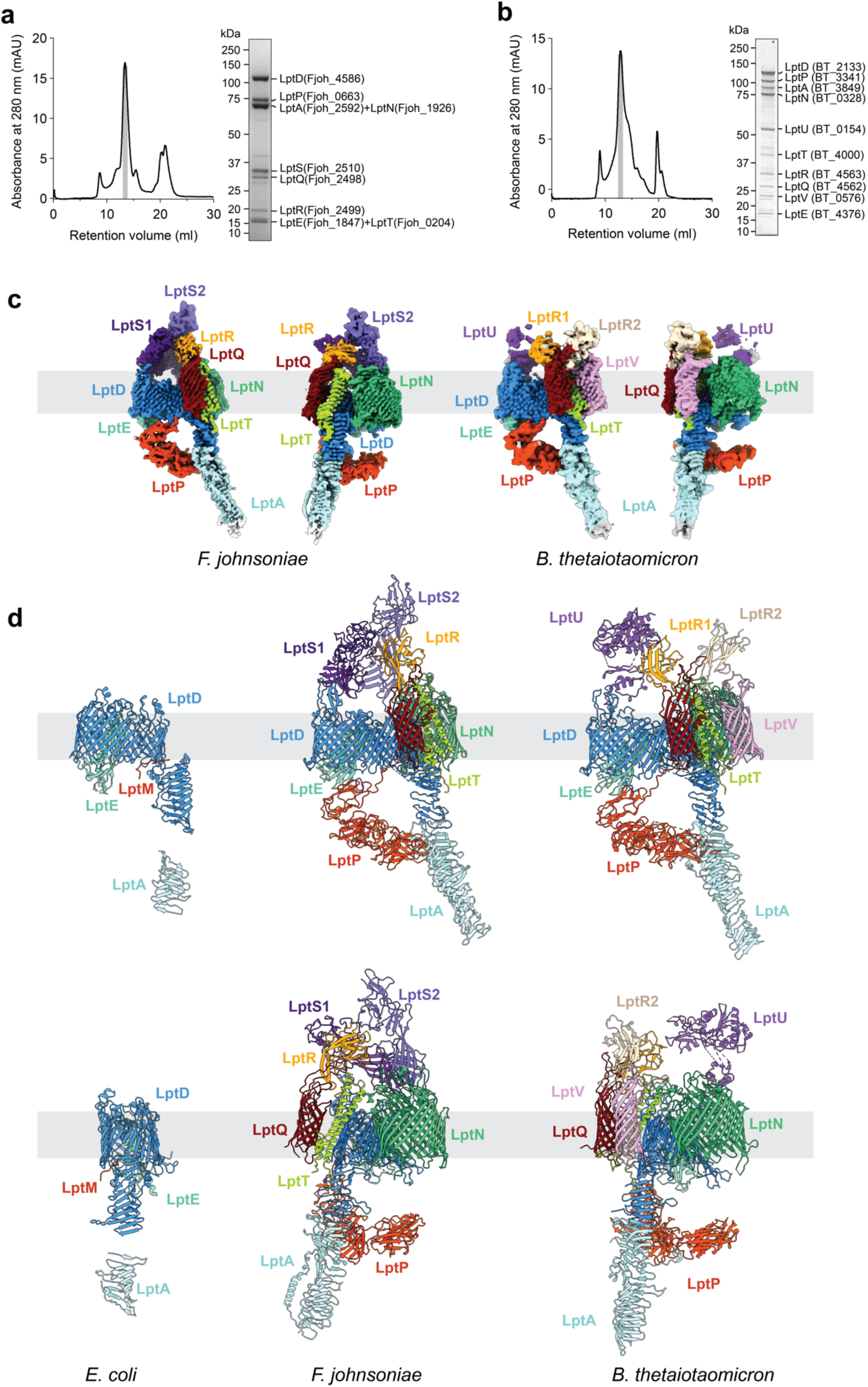
Structures of the endogenous LPS translocons of *F. johnsoniae* and *B. thetaiotaomicron*. **a,b**, Size exclusion chromatography profiles and Coomassie-stained SDS–PAGE gels of the fractions indicated by grey shading for the (**a**) *F. johnsoniae* and (**b**) *B. thetaiotaomicron* LptD preparations. Bands were identified by peptide fingerprinting. Similar data were obtained for three independent preparations. **c,** CryoEM composite volumes for the *F. johnsoniae* and *B. thetaiotaomicron* LPS translocon complexes, with each shown in two views related by a 90° rotation in the plane of the membrane. Unmodelled density is in silver. **d,** Comparison of the *E. coli* (PDB 9KN3), *F. johnsoniae*, and *B. thetaiotaomicron* LPS translocon complexes. The structure of *E. coli* LptA (PDB 2R1A) is also shown for reference. Structures are shown in cartoon representation. The views in the lower row are related to the views in the upper row by a 90° clockwise rotation in the plane of the membrane.

To understand whether similar elaboration of the LPS translocon occurs in other Bacteroidota we also isolated and determined the structure of the endogenous LptD complex (LPT_Bt_) from the major gut commensal *Bacteroides thetaiotaomicron*, a member of the Bacteroidota that is only distantly related to *F. johnsoniae*. Like LPT_Fj_, this complex contains LptA, LptD, LptE, LptN, LptP, LptQ, LptR, and LptT (Fig. 2b). LptS is absent but the complex contains two copies of LptR and two proteins that are not found in the *F. johnsoniae* complex which we name LptU and LptV.

The structures of the LPT_Fj_ and LPT_Bt_ complexes were determined at overall resolutions between 2.9 and 3.3 Å (Fig. 2c,d, Extended Data Figs. 1-4, and Extended Data Table 1) using cryoEM data collected from both native and mildly chemically cross-linked protein samples. Due to their elongated shape the complexes exhibited substantial flexion and so the greatest detail was obtained by using focused refinements over different regions. Atomic models were built and refined in composite volumes as described in the Methods and Extended Data Figs. 1 and 3. Domains distal to the membrane on either side were least well resolved. Refinement of these was constrained by restraint to AlphaFold^33^ models for these domains. O-linked glycans^34^ were built where supported by clear density. LptE, LptP, LptR, LptS, LptU, and LptD_Fj_ are lipoproteins. With the exception of LptD_Fj_, their N-termini are well ordered and suitably positioned for their lipid anchors to insert into the membrane, although no ordered density was observed for the lipid groups themselves. The integral membrane subunits LptD, LptN, LptQ, LptT have long N-terminal periplasmic tails that, with the exception of LptN_Bt_ (see below), are unresolved in the structures. These tails are potentially long enough to reach across the periplasm to interact with the IM LPS transporter.

**Fig. 3:**
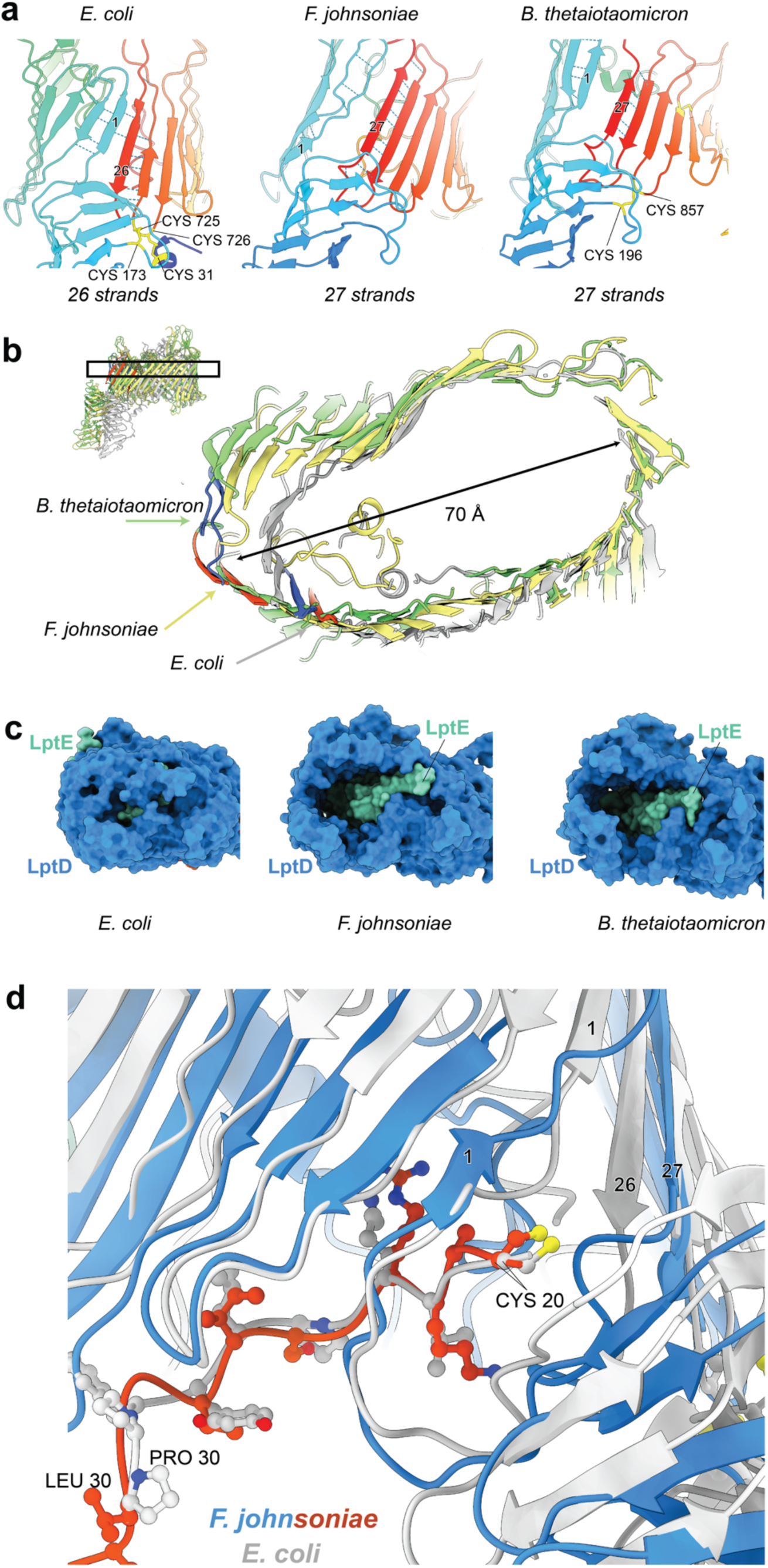
Architecture of the Bacteroidota LptD barrel. **a**, The lateral gate regions of the LptD proteins of *E. coli*, *F. johnsoniae*, and *B. thetaiotaomicron* shown in cartoon representation and rainbow coloured from blue to red. The proteins are aligned via the central portion of the barrel (strands 8-21) and viewed from the direction of the lateral gate. The Bacteroidota proteins have an additional barrel strand and exhibit a positional shift and opening of the lateral gate between the first and last barrel strands, which are numbered. Disulfide bonds present are shown in yellow sticks representation. **b,** A thin slab (as indicated on the inset) through the LptD barrels of *E. coli*, *F. johnsoniae*, and *B. thetaiotaomicron* aligned as in **a**. The proteins are shown in cartoon representation, coloured by species, and are viewed from the extracellular side. The arrows indicate the location of the lateral gate in each protein, which is also marked by colouring the N-terminal strand blue and C-terminal strand red. **c,** The LptD barrels of *E. coli*, *F. johnsoniae*, and *B. thetaiotaomicron* in surface representation viewed from the extracellular side of the membrane. LptE is covered by LptD loops in *E. coli* but not in the Bacteroidota where LptE is exposed and a large cavity is revealed. **d,** Overlay of the LPS translocon complexes of *F. johnsoniae* and *E. coli* (PDB 9KN3) via alignment (RMSD 1.2 Å) of the N-terminus of *F. johnsoniae* LptP (orange) with the N-terminus of *E. coli* LptM (grey), both of which are shown with ball-and-stick sidechains. The LptD proteins are shown in cartoon representation with the N and C-terminal strands numbered to indicated the location of the lateral gate.

Both LPT_Fj_ and LPT_Bt_ complexes contain the LptDE core of the canonical LPS translocon but extensively elaborate this with additional subunits within the OM and on both its periplasmic and extracellular sides (Fig. 2d).

The basic organisation of the LptDE core resembles that of structurally characterised proteobacterial LPS translocons. Thus, LptD forms a large kidney-shaped transmembrane β-barrel plugged at the gate-distal end by the lipoprotein LptE and sealed at the gate-proximal end by LptD extracellular loops, while the periplasmic N-terminus of LptD forms a β-taco domain in front of the barrel lateral gate (Fig. 2d). However, the Bacteroidota LptD barrel contains an additional C-terminal β-strand, giving a total of 27 strands rather than the 26 strands found in previously characterised LptD proteins (Fig. 3a). This is the first example of a single-subunit bacterial OM β-barrel that has an odd number of strands. The odd number of barrel strands changes the organisation of the lateral gate such that the gate strands now interact in a parallel rather than an anti-parallel fashion with both strands running from the periplasm to the cell exterior (Fig. 3a). In both LPT_Fj_ and LPT_Bt_ these strands are separated from each other with no main-chain hydrogen bonds between the strands. Nevertheless, the gate itself remains closed due to interactions between the side chains of the gate strands (Extended Data Fig. 5a). In all previously characterised resting-state LPS translocon structures, the lateral-gate strands are connected by main-chain hydrogen bonds^14^. Whilst the central portion of the LptD barrels (strands 8-21) overlays well between the *E. coli* and *Bacteroidota* proteins (RMSD 1.3 Å), the addition of the extra strand and separation of the gate leads to an expansion of the space enclosed by strands 1-7 and 22-26/27, constituting the gate-proximal part of the barrel (Fig. 3b).

In the proteobacterial LPS translocon three long extracellular loops at the gate-distal end of LptD fold down on top of LptE. This does not happen in the Bacteroidota complexes where the equivalent loops (15-16, 17-18, and 21-22) fold outwards (Fig. 3c and Extended Data Fig. 5b). The result is that there is a substantial cavity on the gate-distal side of LptE that extends from the extracellular environment most of the way across the membrane (Fig. 3c). LptE is now accessible from outside the cell and becomes the sole seal of the transmembrane pore through this part of the LptD barrel.

Previous studies have shown that the N-terminal β-taco domain of LptD docks onto the LptA end of the periplasmic bridge used for LPS transport^35^ and have proposed that it extends the β-taco fold of LptA^7,35,36^. Our structures provide direct structural confirmation of this model by revealing that LptA is assembled onto LptD to form a continuous β-taco structure (Fig. 2c,d). The β-taco in the Bacteroidota LptA proteins has 56 ladder strands, of which 70% can be built in our volumes with the remainder visible at low contour levels (Fig. 2c). This is substantially longer than the 16-stranded *E. coli* LptA protein^36^ (Fig. 2d) likely reflecting the need to span a wider periplasm in the Bacteroidota.

In the LptD protein of *E. coli* and other Enterobacteriaceae the β-taco domain is connected to the C-terminal side of the barrel gate through two disulfide bonds which play an essential role in LptD maturation and function^37,38^. *B. thetaiotaomicron* and most other Bacteroidota conserve the disulfide bond that directly locks the LptD β-taco domain to the periplasmic 24-25 loop of the LptD barrel (Cys196-Cys897 in LptD*_Bt_* corresponding to Cys173-Cys725 in LptD*_Ec_*) (Fig. 3a). However, *F. johnsoniae* LptD contains neither this nor any other disulfide linkages (Fig. 3a).

LptP is a large periplasmic lipoprotein containing five immunoglobulin-like domains. The first domain contacts the rim of the LptD barrel while the remaining four domains form a ring-like arrangement positioned directly beneath the barrel and separated from it by a ∼40 Å gap (Fig. 2c,d and Extended Data Fig. 5c). The ring is closed by interactions between loops at the C-terminal ends of domains 2 and 5 (Extended Data Fig. 5c). The N-terminal end of domain 5 packs against the C-terminal end of the LptA β-taco linking LptP to the periplasmic bridge (Extended Data Fig. 5d). In LPT_Bt_ this interaction is reinforced by contacts with the extended C-terminus of LptA which snakes across the centre of LptP domain 5. This additional interaction is absent in LPT_Fj_ where the equivalent region of LptA packs along the β-taco running in the direction of the inner membrane (Fig 2d).

The N-terminal region of the LptP polypeptide forms an extended strand that runs along the inner face of the periplasmic rim of the LptD barrel before exiting through the lower part of the lateral gate (Figs. 2d and 3d). This places the lipidated N-terminal cysteine immediately outside the gate, where its unresolved acyl chains would insert into the inner leaflet of the membrane. Remarkably, the N-terminal polypeptide of LptP occupies the same position as the lipidated N-terminal peptide of the small LptM subunit found in enterobacterial LPS translocons^12,14–16^ (Fig. 3d) despite the absence of detectable sequence similarity between the two proteins. This striking structural correspondence suggests that the N-terminal region of LptP may perform functions equivalent to those of LptM. By contrast, LptP is structurally unrelated to LptY (formerly YedD), the other periplasmic subunit of the *E. coli* LPS translocon^14,17^, nor does it occupy an equivalent space under LptD.

The Bacteroidota-specific transmembrane subunits form two sub-complexes that bind externally to LptD on either side of the lateral gate (Fig. 2d and 4a). The C-terminal side sub-complex is formed solely by LptN. The N-terminal side sub-complex is organised by LptT which associates directly to LptD. LptT then acts as the primary binding site for a complex between the membrane protein LptQ and the SLP LptR, and in *B. thetaiotaomicron* for LptV. On the periplasmic side of the membrane the two sub-complexes bind to opposite sides of the LptD β-taco domain, while at the extracellular side of the membrane the sub-complexes are in direct contact. Opening of the lateral gate will be linked to corresponding movements of the sub-complexes attached to either side of the gate (or vice versa). Thus, opening the lateral gate will also break the extracellular contact between the sub-complexes that would otherwise block substrate release through the lateral gate.

**Fig. 4:**
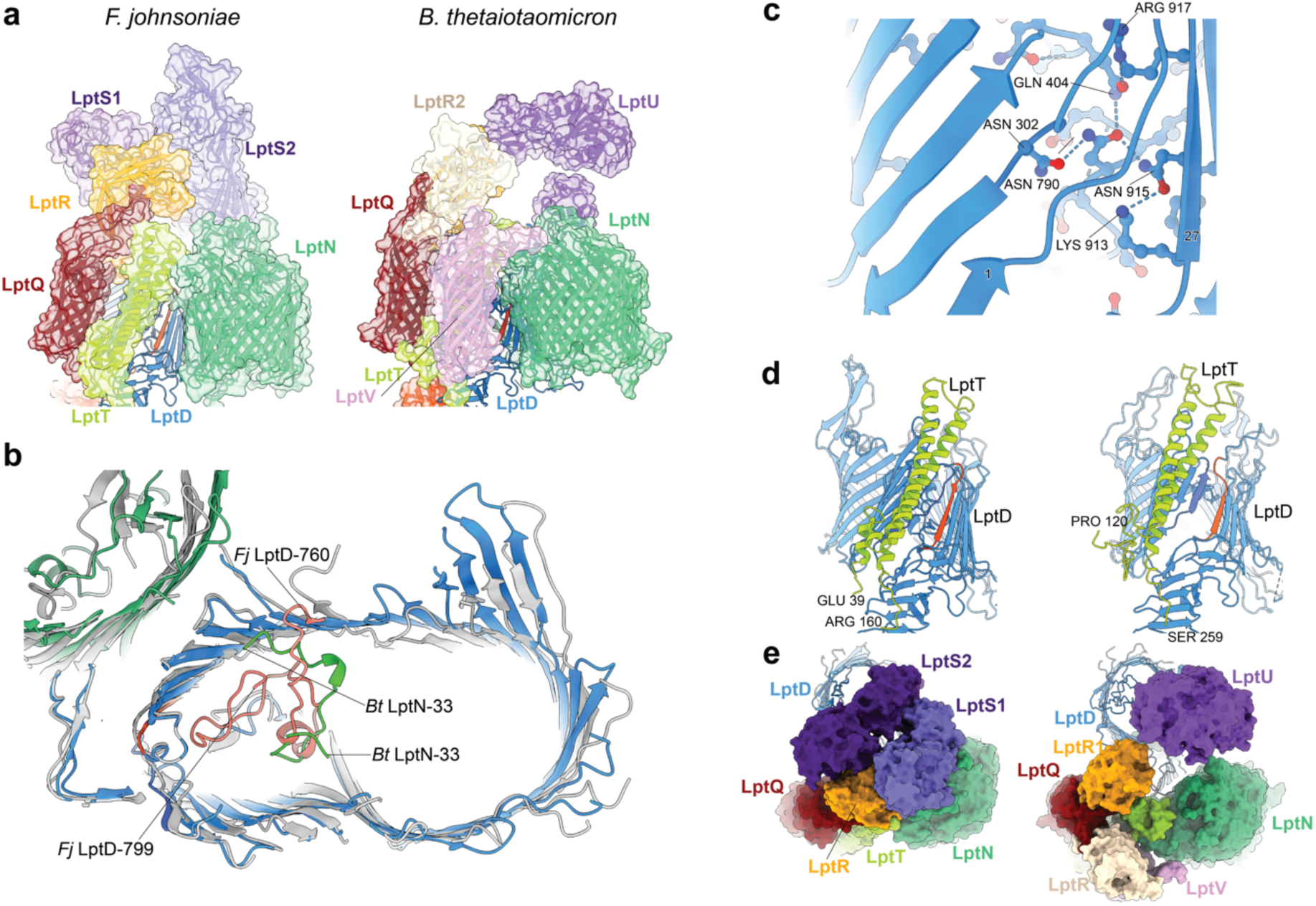
Further structural analysis of the Bacteroidota LPS translocons. **a**, The Bacteroidota LPS translocon complexes viewed from in front of the lateral gate of LptD. Subunits around the lateral gate are show in cartoons representation with semi-transparent surfaces. LptD is shown in cartoons representation with the strand on the C-terminal side of the lateral gate in red. **b,** In the *F. johnsoniae* (*Fj*) LPS translocon (grey) the LptD loop between strands 20 and 21 (salmon) packs within the LptD barrel. By contrast, the N-terminus of LptN (green) occupies the same region of the barrel (blue) in the *B. thetaiotaomicron* (*Bt*) translocon. Only LptD and LpN are depicted. The proteins are viewed from the periplasm and shown in cartoons representation. The structures are aligned across the entire LptD barrel (RMSD 1.3 Å) and sliced to remove their extracellular sides. **c,** In *F. johnsoniae* LptD, Asn790 in the periplasmic loop between strands 20 and 21 forms a hydrogen-bonding network that ties together both sides of the lateral gate (through Asn302 and Asn915) and extracellular loop 7/8 (through Gln404) that seals the gate-proximal pore. Cartoons representation with ball and stick side chains. The lateral gate strands are numbered. **d,** The helical LptT protein packs at the LptD lateral gate. The proteins are shown in cartoons representation with the LptD gate strands highlighted blue at the N-side and red at the C-side. **e,** The Bacteroidota LPS translocon complexes viewed from outside the cell. LptD is shown in cartoons representation and the remaining subunits in surface representation. LptA, LptE and LptP are omitted for clarity.

LptN is a 22-stranded OMP of oval cross-section (Extended Data Fig. 6a-c), structurally unrelated to other characterized 22-stranded OMPs including the SusC protein family. The interior of LptN is closed to the extracellular environment by interactions between the extracellular inter-strand loops (Extended Data Fig. 6b,c). The central cavity is largely occupied by a structured region of ∼100 residues preceding the first β-strand (Extended Data Fig. 6c). In LPT_Fj_ the N-terminal tail of LptN becomes disordered once it emerges from this cavity. By contrast, in LPT_Bt_ part of the N-terminal tail extends into the LptD barrel and packs against LptE (Fig. 4b). The equivalent region of the LptD cavity is occupied in LPT_Fj_ by a partially ordered periplasmic loop from LptD (between β-strands 20 and 21) (Fig. 4b,c) and in the enterobacterial LPS translocon by the C-terminal part of the LptM subunit^12^.

The first three β-strands of LptN pack against the final four β-strands of LptD, with additional contacts made by the inter-strand loops surrounding these regions. LptN also binds to the β-taco domain of LptD through interactions with the two final β-taco loops. This helps tie the β-taco bridge to the C-terminal side of the LptD barrel, a role performed by the disulfide bonds in *E. coli* LptD. Taken together these interfaces bury a surface of 1092 Å^2^ in LPT_Fj_ and 2242 Å^2^ in LPT_Bt_. Even the smaller contact area in LPT_Fj_ is equivalent to those of many high affinity antibody-antigen complexes, implying that a very stable association exists between LptN and LptD in both species and that LptN will move with the C-terminal part of the LptD barrel.

LptN also interacts with the extracellular end of the LptQRT(V) subcomplex (Fig. 4a). In LPT_Fj_ this interaction is mediated solely by contact between the 5-6 loop of LptN and LptT, burying a surface of 250 Å^2^. In LPT_Bt_, by contrast, the LptN 5-6 loop makes more extensive contacts with LptT that are augmented by interactions from the LptN strand 1-2 loop, burying a total of 703 Å^2^ between the chains, and a small contact with LptV burying another 80 Å^2^.

LptT is at the core of the subcomplex on the N-side of the LptD gate. LptT is a highly unusual OM protein that is constructed from membrane-integrated α-helices rather than forming a β-barrel (Fig. 4d). Only a few other bacterial OM proteins with transmembrane α-helices are known^39–42^ and unlike LptT these proteins all form homo-oligomers in the membrane. LptT comprises two long transmembrane helices (TMHs) that cross the OM and protrude into the extracellular space, and two short hydrophobic helices that insert part way across the membrane from the periplasmic side and pack against the transmembrane helix pair. LptT is oriented approximately parallel to the N-terminal strands of the LptD barrel and so tilted ∼30 ° from the membrane normal when viewed towards the lateral gate. LptT binds the LptD barrel through interactions that the extracellular end of its C-terminal TMH makes with barrel strands 1 and 2, and with loops 1-2, 3-4, and 5-6. The periplasmic end of the same TMH, together with the following loop, pack against the top of the LptD β-taco. Together, these interactions between LptT and LptD bury ∼1,200 Å^2^ of surface implying a stable association that will cause the LptT-centred sub-complex to move with the N-terminal part of the LptD barrel.

LptQ is an 8-stranded β-barrel OMP. An unusual feature of the barrel is that the β-sheet is interrupted in the extracellular half of the membrane by strand 1 transitioning into an amphipathic helix (Extended Data Fig. 6d). The extracellular loops of LptQ anchor one copy of the SLP subunit LptR in Lpt_Fj_ and two copies in Lpt_Bt_, with the second copy of LptR in Lpt_Bt_ interacting with a large insertion in the loop between LptQ strands 5 and 6 (Fig. 4a,e and Extended Data Fig. 6d). In Lpt_Bt_ the LptQR_2_ unit is attached to the periplasmic end of LptT through its short N-terminal helix burying ∼730 Å^2^ of surface. This positions the LptQR_2_ unit close to, but not in contact with, LptD. By contrast, the LptQR unit in the Lpt_Fj_ complex makes additional interactions at the extracellular side of the membrane with LptT that bring it into contact with the strand 5-6 loop of the LptD barrel.

Lpt_Bt_ contains an additional 8-stranded β-barrel subunit, LptV, which further extends the architecture of the subcomplex at the N-side of the lateral gate. LptV packs against the side of LptT opposite from LptQ through interactions with the N-terminal TMH of LptT (Fig. 4a,e). LptV is also held in place by interactions with LptN, as described above, and with the second copy of LptR present in the Lpt_Bt_ complex.

On the extracellular side of the complex are a collection of SLP subunits that form a partial dome above the LptD barrel (Figs. 2c,d and 4a,e). These include the previously described LptR protein found in both LPT_Fj_ and LPT_Bt_. The remaining SLPs differ between the two complexes with LPT_Fj_ having two copies of LptS and LPT_Bt_ a single LptU subunit.

In LPT_Fj_ the two LptS subunits, each of which is composed of two β-strand domains, form a head-to-head dimer related by a 180 ° rotation along the axis between them (Extended Data Fig. 6e). This arrangement allows the two halves of the homodimer to make distinct interactions with the underlying translocon complex. The N-terminal domains of the two subunits anchor the dimer to the gate-distal end of the complex through separate interactions with LptD and LptN. LptS copy 1 contacts extracellular loop 19-20 in LptD, while stands β7 and β8 of LptS copy 2 form an intermolecular β-sheet with the 9-10 extracellular loop of LptN. From these anchor points, the LptS dimer extends across the translocon complex, with the C-terminal domains of the two subunits making different interactions with the top of LptR (Fig. 4e).

The LptU subunit in LPT_Bt_ is a three-domain protein. It has a Peptidase S41 fold but lacks the catalytic Ser-Lys pair^43^ and is therefore unlikely to function as a protease. LptU occupies a similar position to LptS1 in the LPT_Fj_ complex with the N-terminal domain binding to LptD loops 19-20 and 21-22. The two remaining domains are only weakly visible in the EM volume. Although they are directed towards LptR1 they do not touch it (Fig. 4e).

### Trajectory of translocon opening

Although the overall organisation of the gate-adjacent subunits is conserved between LPT_Fj_ and LPT_Bt_ their precise positioning varies in a systematic way between the two complexes in a way that appears to mimic the first stages in translocon opening. These differences are most obvious when the two structures are aligned on the gate-distal end of the LptD barrel (Fig. 5a). Although the lateral gate is still closed in both complexes, LPT_Bt_ has a series of linked conformational differences from LPT_Fj_ that would be consistent with a state that is further along an incipient opening trajectory.

**Fig. 5:**
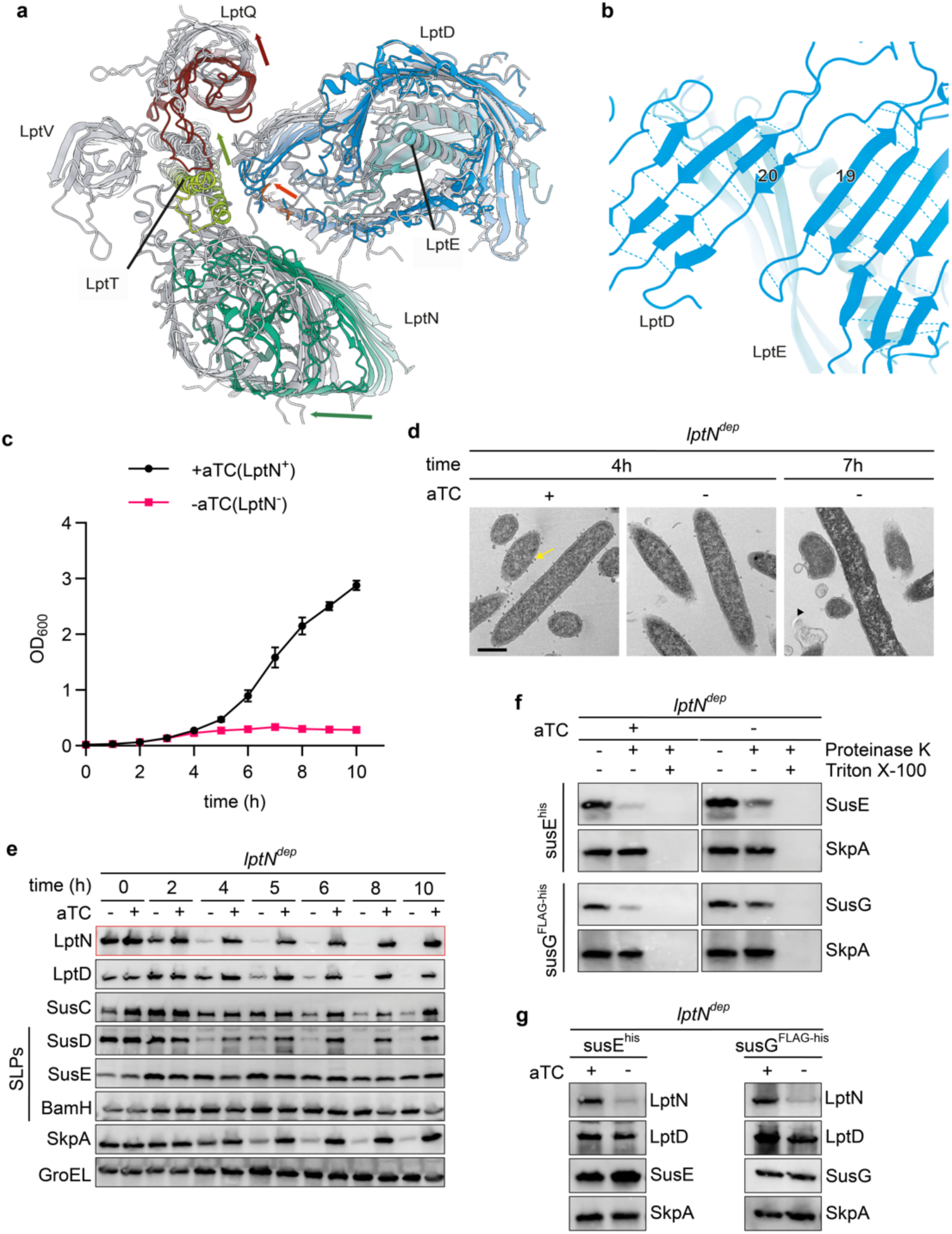
Trajectory of translocon opening and demonstration that LptN is an essential protein in *F. johnsoniae*. **a**, Conformational differences between the intra-membrane components of the *Flavobacterium johnsoniae* (coloured by subunit) and *Bacteroidota thetaiotaomicron* (silver) complexes demonstrated by aligning on strands 8-21 of the LptD barrel. The proteins are shown in cartoons representation and and sliced to remove their periplasmic sides. The C-terminal strand of the LptD barrel is shown in red to highlight the location of the lateral gate. Arrows highlight subunit and gate movements between the two complexes. **b,** The *B. thetaiotaomicron* LptD barrel contains a break in the β-sheet between strands 19 and 20. **c-g,** The LptN depletion strain (*lptN*^dep^) was cultured in rich (CYE) medium. In the -aTC samples the aTC inducer was removed at 0 h to initiate LptN depletion. SkpA, is a periplasmic protein; GroEL a cytoplasmic protein. Similar data were obtained for three biological repeats. **c,** Growth curves (means ± 1 SD, for four biological repeats). **d,** Whole cell immunoblots. SLPs, surface lipoproteins. **e,** Representative transmission electron microscopy images during LptN depletion. Yellow arrows highlight budding OM vesicles, black arrows highlight OM blebbing. Scale bar, 500 nm. **f,** Depletion of LptN results in intracellular accumulation of the SLPs SusE and SusG as assessed by Proteinase K accessibility. Triton X-100 permeabilises the OM. Reactions were analysed by immunoblotting. **g,** Whole cell immunoblots of the samples used in **d** showing that LptD is not appreciably depleted.

The LptD barrel undergoes a very unusual structural change in the LPT_Bt_ complex involving β-strands 19 and 20 which separate and twist inwards (Fig. 5b). Despite this break in the β-sheet, the barrel cavity remains isolated from the surrounding membrane due to interactions between bulky side chains. The strand separation creates the equivalent of a one-strand gap relative to the LptD_Fj_ barrel. This shifts the register of the subsequent strands by one position towards the C-terminus, effectively expanding the barrel by one strand (Figs. 3b and 5a). As a consequence, the lateral gate is rotated clockwise when viewed from outside the cell, such that the C-terminal strand of the LptD_Bt_ gate occupies an equivalent position to the N-terminal strand of the LptD_Fj_ gate (Figs. 3b and 5a). This change is accommodated by an outward movement of β-strands 1–6, particularly at the extracellular side of the membrane, that widens both the gate-proximal pore and the lateral gate itself (Figs. 3a,b and 5a).

LptN is bound to C-terminal side of the LptD lateral gate, so the clockwise movement of this part of the LptD barrel in LPT_Bt_ also moves LptN clockwise (Fig. 5a). As a consequence, the gap between LptD and LptN is wider in LptD_Bt_ relative to LptD_Fj_ (Fig. 5a).

On the other side of the lateral gate LptT moves with the N-side of the gate while remaining connected extracellularly with LptN as LptN moves with the C-side of the gate (Fig. 5a). LptT therefore mechanically couples movements on the opposite sides of the lateral gate. Notably, these conformational changes actually bring LptT and LptN into closer contact, and so strengthen rather than open the interface between the gate-adjacent subcomplexes. Movement of LptT, in turn, carries the associated LptQ, LptR, and LptV subunits with it. This results in the LptQR1R2 subcomplex tilting away from the extracellular side of the LptD barrel even though this part of the barrel has expanded outwards. From this position LptR1 is unable to interact with LptU in a manner analogous to the LptR interaction with LptS1 and LptS2 in LPT_Fj_ (Fig. 4e).

A further difference between the two structures is that in LPT_Fj_ the LptD periplasmic loop between β-strands 20 and 21 is inserted into the gate-proximal end of the LptD pore and this loop is used to connect the elements that surround the lateral gate (Figs. 4b,c). Asn790 within the loop forms hydrogen bonds with Asn302 in β-strand 2 at the N-terminal side of the lateral gate, with Asn915 in β-strand 27 at the C-terminal side of the lateral gate, and with Gln404 in the β7–β8 extracellular loop that seals the gate-proximal end of the barrel (Fig. 4c). The effect is to couple these parts of the structure together. No equivalent interactions are present in LPT_Bt_ where the barrel contains the N-tail of LptN rather than the LptD 20-21 periplasmic loop.

Thus, there is a complex change in morphology between the two complexes driven by a web of interconnecting structural changes between interacting subunits. The structures do not establish which of these structural changes initiate this transition.

### Subunit conservation

Analysis of the occurrence of the LPT*_Fj_* and LPT*_Bt_* subunits across the Bacteroidota suggest that LptA, LptD, LptE, LptN, LptP, and LptT constitute the invariant core of the Bacteroidota LPS translocon (analysis of representative species is presented in Extended Data Table 2, Supplementary Information Table S1). Although LptQ and LptR are also normally present, they are sporadically missing from species across multiple Bacteroidota Classes. The remaining subunits of LPT_Fj_ and LPT_Bt_ are not widely conserved. The genes encoding the Bacteroidota LPS translocon are genetically unlinked, with the exception that LptQ and LptR are universally coded by adjacent genes consistent with their role in forming a structural subcomplex within the translocon. LptP is also conserved across the seven additional phyla of the wider FCB (Fibrobacterota–Chlorobiota–Bacteroidota) superphylum, with LptT present in six of these, suggesting that these phyla also possess expanded LPS translocons that share features of the Bacteroidota system (Extended Data Table 2).

### LptN is an essential protein and involved in SLP export

To investigate the function of the additional subunits of the Bacteroidota LPS translocon, we attempted to delete their coding genes from *F. johnsoniae*. We also targeted a gene (*fjoh_1207*) coding for a LptT paralogue (62% amino acid identity) which we designate LptT2. LptT2 is expressed at much lower levels than LptT under the growth conditions used here^44^ but was detected at low abundance in the LPT_Fj_ preparation by peptide mass spectrometry.

We were unable to delete the gene encoding LptN. Likewise, we were unable to generate a double mutant lacking both LptT and LptT2 even though their coding genes could be individually deleted. These observations suggest that LptN is essential, as is the presence of at least one LptT paralogue. Published whole-genome mutagenesis studies in *B. thetaiotaomicron* indicate a similar pattern, with LptA, LptD, LptE, LptN, and LptT being essential, whereas mutants lacking any of the remaining LPT_Bt_ subunits still being viable^45–47^.

We confirmed that LptN was essential in *F. johnsoniae* using the same conditional depletion strategy employed for LptD (Fig. 5c). Upon prolonged LptN depletion we observed that LptD was also lost indicating that LptN is required for the long-term stability of LptD (Fig. 5e). Since LptD is itself essential, the eventual loss of LptD would be sufficient to account for the lethality of LptN depletion. Nevertheless, cell growth slowed before any appreciable loss of LptD, indicating that LptN depletion has deleterious effects independent of its effect on LptD levels. The dependence of LptD on LptN was not reciprocal, as LptN remained stable following LptD depletion (Fig. 1b,e). This excludes secondary loss of LptN as an explanation for the effects of LptD depletion on SLP biogenesis in our earlier experiments. Prolonged LptN depletion resulted in severe disruption of membrane organisation (Fig. 5d).

To assess the role of LptN in SLP export without the confounding effect of LptD destabilisation, we examined early depletion time points when LptN was almost completely depleted but LptD levels remained comparable to those in non-depleted cells. At such time points loss of SusD is evident (Fig. 5e) and protease protection assays show accumulation of the tested SLPs inside the cell (Fig. 5f,g). From these results we conclude that LptN is involved in SLP export in *F. johnsoniae*.

### Phenotypic effects of loss of non-essential LPT*_Fj_* components

We succeeded in generating individual *F. johnsoniae* mutants lacking LptP, LptQ, LptR, LptS, LptT, or LptT2. Immunoblotting with available subunit-specific antibodies confirmed that only the targeted subunit was absent from the deletion strains (Extended Data Fig. 7a). Isolation of LptD complexes from these strains indicated that all subunits other than the target subunit were still present, although in some cases at reduced levels suggesting effects on complex stability (Extended Data Fig. 7b,c). Exceptionally, removal of LptQ prevented LptR incorporation into the complex consistent with the structural role of LptQ in anchoring LptR.

For the strains lacking LptP, LptQ, or LptR we observed modest reductions in growth rate in rich medium (Extended Data Fig. 7d). Fewer OMVs were formed by these strains but otherwise there was no change in cell morphology (Extended Data Fig. 7e). All three of these mutants showed greatly enhanced sensitivity to the detergent SDS, indicating defects in OM function (Fig. 6a). The *lptP* mutant also exhibited a very large increase in sensitivity to the high molecular weight antibiotic vancomycin suggesting an increase in OM permeability in this strain (Fig. 6a). Nevertheless, the OM of this mutant remained intact, as judged by cell imaging (Extended Data Fig. 7e) and by retention of the periplasmic protein SkpA in the cells (Extended Data Fig. 7a). The *lptS*, *lptT*, and *lptT2* mutants showed no detectable changes in growth rate in rich medium, cell morphology, or OM function relative to the parental strain (Fig. 6a and Extended Data Fig. 7d,e).

**Fig. 6:**
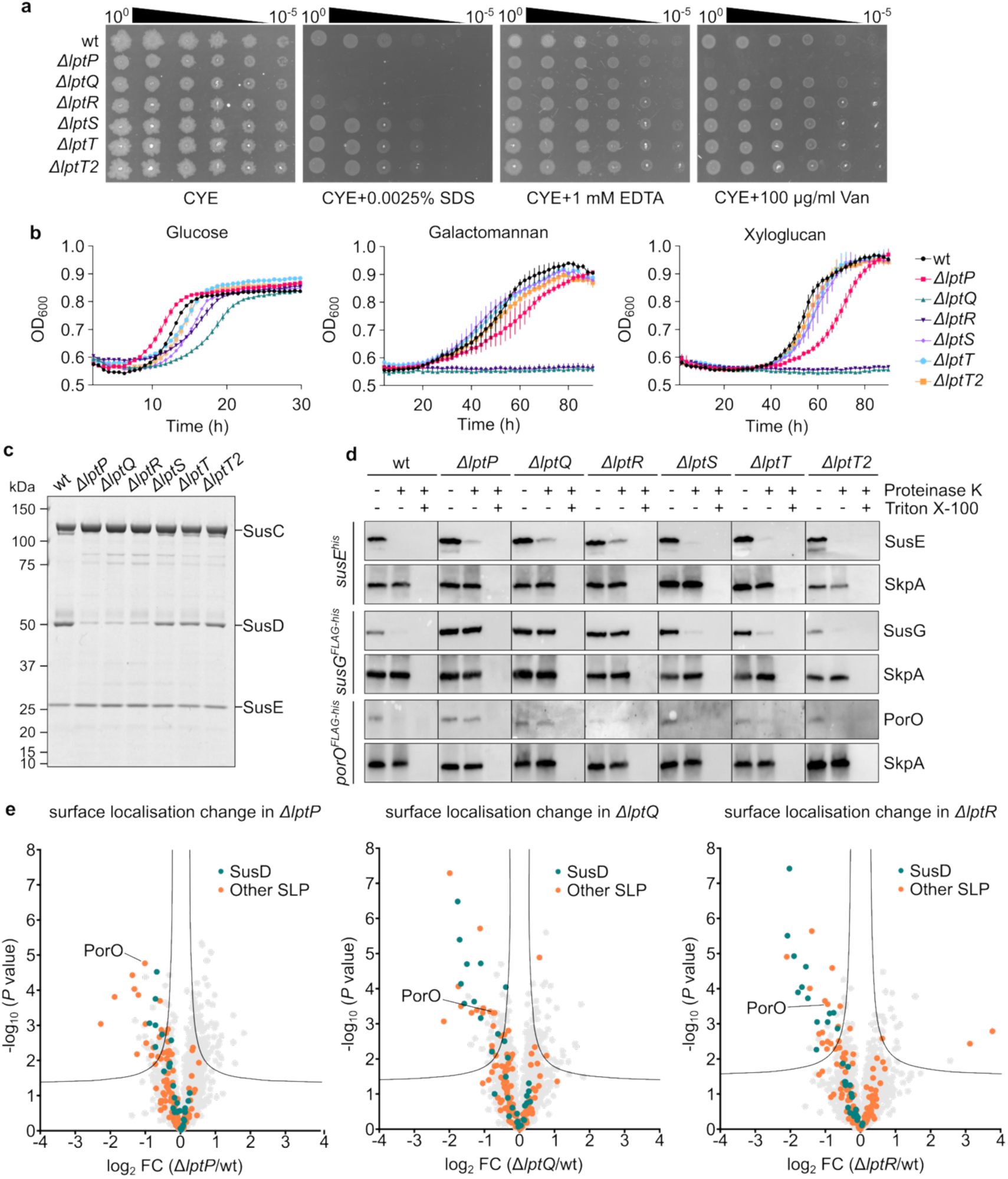
Phenotypic effects of removing non-essential LPT_Fj_ subunits. **a**, OM integrity assays. Cells were grown on CYE agar with the indicated additions. Van, vancomycin. **b,** Growth curves on minimal medium containing either glucose or galactomannan or xyloglucan as sole carbon source. Shown are the means ± 1 SD from three technical repeats. **c,** Native SusC complexes affinity-purified via a N-terminal Twin-Strep tag on SusC. Coomassie-stained SDS–PAGE gel. **d,** Surface exposure of the SLPs SusE, SusG, and PorO assessed by Proteinase K accessibility. Triton X-100 permeabilises the OM. Reactions were analysed by immunoblotting. SkpA, is a periplasmic protein. Corresponding whole cell immunoblots in Extended Data Fig. Xg. **e,** Identification of cell surface proteins that re-localise to the cell interior in mutants lacking LPT_Fj_ subunits. Cell surface exposure was quantified by biotinylation with the poorly OM-permeable reagent sulfo-NHS-LC-LC-biotin. Comparative analysis of biotinylation in the mutant versus the wild type strain after correction for changes in whole cell protein abundance. FC, fold change. A significance threshold is drawn according to a two-tailed t-test with a permutation-based false discovery rate of 0.05 and a S_0_ of 0.1. Data are from four biological repeats. wt, wild type. Similar data were obtained for three (**a,b,d**) or two (**c**) biological repeats.

We used multiple methods to assess whether the subunit-deleted strains had defects in SLP export. Export defects were consistently observed in mutants lacking LptP, LptQ, or LptR, but not in the other deletion strains.

We first assessed growth of the mutants on the polysaccharides galactomannan and xyloglucan which both require SLP-containing systems for breakdown and uptake^48^. Strains lacking LptQ or LptR failed to grow on either polysaccharide, while the LptP-deficient strain exhibited delayed growth (Fig. 6b). These observations are consistent with these subunits having a role in SLP biogenesis. Control experiments showed that all three strains showed only modest at most growth defects on glucose, a saccharide that can be transported into cells without the involvement of SLPs (Fig. 6b).

We next examined the biogenesis of the SusD and SusE SLP subunits of the major *F. johnsoniae* SUS complex. Cellular levels of SusD, but not SusE, were substantially reduced in the *lptP*, *lptQ*, and *lptR* mutants (Extended Data Fig. 7a). Consistent with these observations, isolation of SusC complexes from these mutants showed that they had selectively lost most of their SusD subunits while still efficiently incorporating the second SLP subunit SusE (Fig. 6c). These observations suggest that LptP, LptQ, and LptR are involved in the biogenesis of SusD but are not important for the export of SusE.

The compositions of the LptD complexes isolated from the deletion strains provide insight into the biogenesis of the SLP subunits of the LPS translocon (Extended Data Fig. 7b,c). The translocon SLP LptS remained associated with LptD in strains lacking LptP, LptQ, LptR, or LptT, demonstrating that none of these subunits is required for LptS export. Similarly, the translocon SLP LptR remained incorporated into LptD complexes in the absence of LptP, indicating that LptP is dispensable for LptR export.

We next used protease protection assays to ask whether representative SLPs could still reach the cell surface in the mutant strains (Fig. 6d and Extended Data Fig. 7f). SusE export was only weakly affected at most by any of the deletions, in agreement with the SusC complex purification data. By contrast, export of SusG and PorO was completely blocked in the Δ*lptP*, Δ*lptQ*, and Δ*lptR* mutants, with the unexported proteins accumulating inside the cells. Thus, the export of some SLPs is entirely dependent on a subset of the novel subunits in the *F. johnsoniae* LPS translocon.

Taken together these experiments establish that LptP, LptQ, and LptR are involved in SLP export with individual SLPs exhibiting either a very high (SusD, SusG, and PorO) or very low (SusE, LptS, LptR) dependence on these subunits. Cells unable to export the LptPQR-dependent SLPs remain viable, suggesting that the non-transported SLPs do not block the pathway through which LPS moves through the translocon.

We sought to generalise these findings by assessing the effects of the loss of LptP, LptQ, and LptR on SLP export across the proteome. To do this we treated cells with sulfo-NHS-LC-LC-biotin which adds biotin labels to those proteins with which it reacts. Because the OM is only sparingly permeable to this reagent, any SLPs that fail to be transported from the periplasm to the cell surface due will exhibit reduced biotinylation. We validated this methodology with test SLPs that have differing responses to Lpt subunit depletion (Extended Data Fig. 8a,b). We then applied this analysis to the whole proteome using label-free quantitative mass spectrometry, correcting for any differences in the cellular levels of each protein between the parental and mutant strains (Fig. 6e). We found that the majority of proteins with significantly reduced biotinylation in the Lpt mutant strains are SLPs (Fig. 6e, Extended Data Fig. 8c and Extended Data Table 3). Thus, the conclusion that LptP, LptQ, and LptR are involved in SLP export is substantiated at the whole proteome level.

There was a strong overlap between the SLPs mislocalised in all the mutant strains (Extended Data Fig. 8d and Extended Data Table 3). The highest correlation between the patterns of changes in SLP surface exposure was between the LptQ- and LptR-deficient strains as expected for two proteins that form a subcomplex within the LPS translocon (Extended Data Fig. 9). Notably these strains showed greater depletion of SusD family proteins in the LptP-deficient mutant (Fig. 6e, Extended Data Fig. 9). There was no correlation between mislocalisation and protein size, and no common feature of the SLPs was found to be associated with the degree of mislocalisation.

## Discussion

Here we identify the LPS translocon as the long-sought SLP exporter in the Bacteroidota. We show that this translocon is highly elaborated beyond the canonical LptADE core, with at least 7 additional subunits in the two examples characterised here. Four of these additional subunits are shown to be involved in SLP export (Fig. 7a) suggesting that the expansion of the LPS translocon has been driven, at least in part, by the requirements of SLP translocation. We find that two of the additional subunits, LptN and LptT, which are conserved across the Bacteroidota, are the products of essential genes. Thus, in the Bacteroidota the LPS translocon has an essential core that has been increased by two subunits relative to the canonical LPS translocon.

**Fig. 7:**
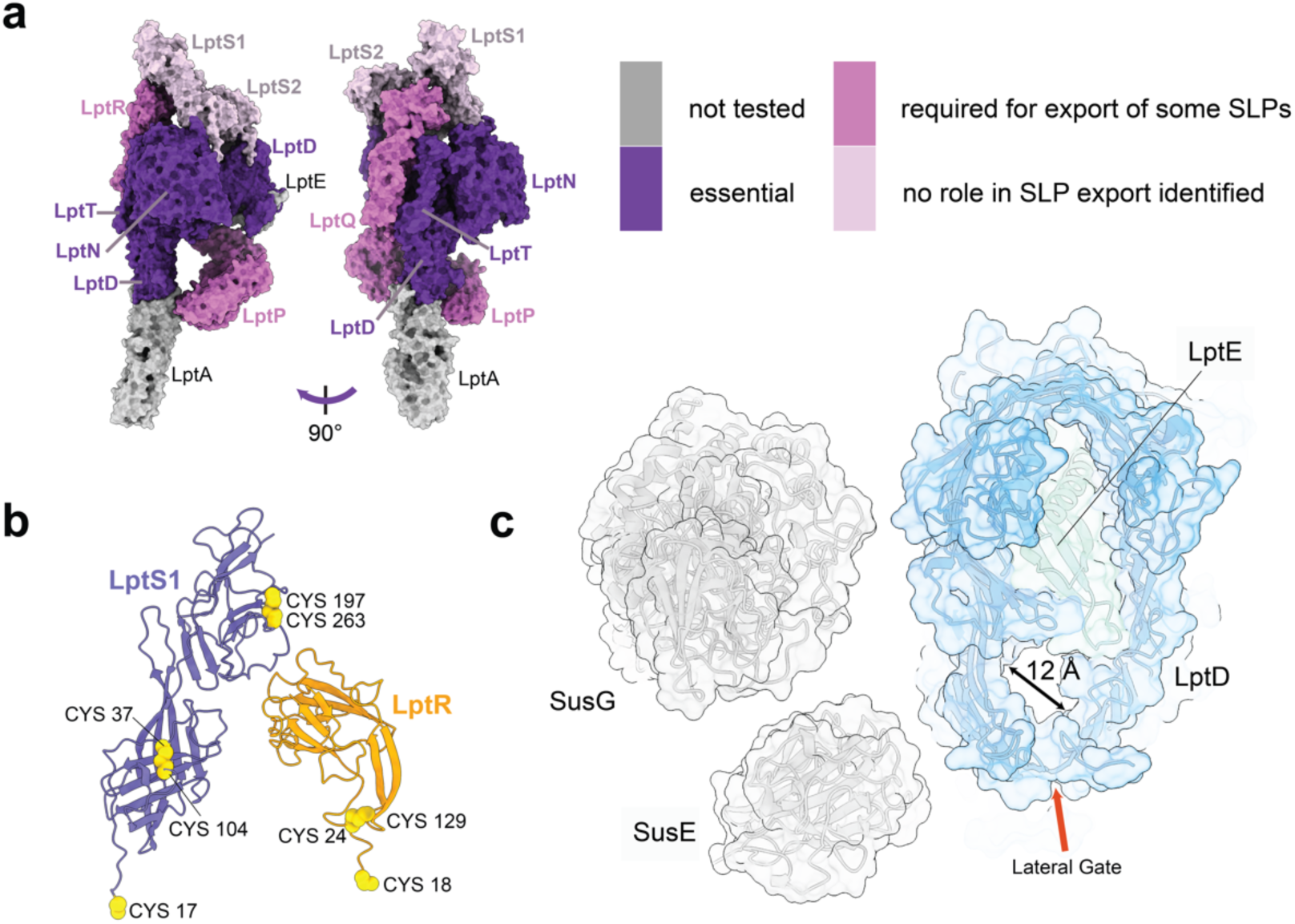
Implication of the Bacteroidota LPS translocon structures for SLP export. **a**, The *F. johnsoniae* translocon shown in a surface representation with subunits coloured by phenotype after deletion or depletion. **b,** The LptR and LptS components of the LPS translocon are SLPs that must be exported after insertion of disulfide bonds. F. johnsoniae proteins LptR and LptS1 are shown in cartoon representation with the N-terminal cysteine that forms the site of lipidation and the intra-domain disulfide bonds shown in yellow spacefilling representation. **c,** Size comparison between the LptD barrel and LptE plug complex from *B. thetaiotaomicron* and two experimentally determined *F. johnsoniae* SLP substrates (SusE and SusG). The proteins are shown as cartoon representations within semi-transparent surfaces. The LptD extracellular loop between β7–β8 has been removed to help illustrate the size of the LptD pore. Very extensive rearrangement of LptD would be required to accommodate even the smaller SusE substrate.

The expansion of the LPS translocon in the Bacteroidota has a striking parallel in the BAM system for OM protein insertion which is also extensively elaborated in the Bacteroidota relative to the paradigm in other phyla^20,21^. The expansion in both cases includes a partial extracellular canopy composed of SLPs and an increase in the number of essential membrane subunits.

Our structural and functional data place important constraints on the mechanism of SLP export by the Bacteroidota LPS translocon.

First, the shared use of the LPS translocon for SLP and LPS export suggests that both substrates employ the same underlying transport mechanism. This expectation is reinforced by the experimental demonstration that SLP export requires the LptD protein that is central to LPS transport (Fig. 1d), and by the grossly similar amphipathic structures of the two types of substrates. A consequence of this shared mechanism is that the adaptations enabling SLP export will also impact LPS transport. For example, the extracellular interaction between the subcomplexes flanking the LptD lateral gate (Fig. 4a,e) would block exit of LPS from the LptD barrel, and so this interface needs to be opened during LPS transport.

Second, the Bacteroidota-specific subunits of the translocon, including those experimentally implicated in SLP export, cluster around the LptD lateral gate. Their location implies that the mechanistic modifications that allow the LPS translocon to export SLPs involve modulating lateral gate function.

Third, SLPs differ markedly in their dependence on the non-canonical LptP, LptQ, and LptR subunits for export (Figs. 6c–e and 7a). This suggests that LptPQR-dependent SLPs may require additional assistance in their export and raises the possibility that different classes of SLP substrate may be handled differently by the translocon.

Fourth, at least some SLPs are likely exported in a folded state. We base this deduction on our structural data which show that the LptR and LptS SLP subunits of the translocon contain disulfide bonds (Fig. 7b). Because disulfide bonds are inserted into proteins in the periplasm^49^ this indicates that these SLPs are at least partly folded prior to export.

Fifth, the ∼25 Å diameter open state of the LptD pore that is seen in the proteobacterial LPS translocon^12^, while appropriate for LPS transport, is far too narrow to accommodate the globular domains of folded SLPs (Fig. 7c). One mechanistic solution to this problem would be for most SLPs to be transported in an unfolded or substantially unfolded state, despite the just-presented evidence that some SLPs are exported at least partly folded. In this model the non-canonical subunits might help maintain the substrate in a transport-competent state and assist in folding at the cell surface following transport. Alternatively, the Bacteroidota LPS translocon could undergo a much larger opening than the canonical translocon, with the Bacteroidota-specific subunits facilitating formation of this expanded state. The gate-proximal part of the Bacteroidota LptD barrel is already somewhat wider than in proteobacterial LptD as a consequence of the additional barrel strand, further barrel expansion through strand separation, and the partially open resting state of the lateral gate (Figs. 3a,b and 5a,b). Nevertheless, it seems unlikely that expansion of the gate-proximal pore alone could provide an export route for the largest SLP substrates which are up to ∼50 Å in diameter (Fig. 7c). We therefore propose that SLP export involves displacement of the LptE plug from the LptD barrel in order to open up the whole barrel for SLP transport. In the proteobacterial translocon LptE remains tightly associated with the LptD barrel even after opening of the gate-proximal pore^12^ due to interactions with the surrounding barrel walls and with extracellular loops that pack onto LptE. However, the extracellular loop interactions are absent in the Bacteroidota translocon (Fig. 3c and Extended Data Fig. 5b). We propose that a larger-than-canonical expansion of the gate-proximal region, facilitated by the additional translocon subunits, could disrupt the remaining interactions between LptE and the barrel wall through a concomitant small expansion of the gate-distal end of the pore.

Since SLPs use the LPS translocon for export across the OM, it is most likely that they reach the translocon by the same route as LPS by travelling through the LptA bridge. This model would explain previous observations that SLP export in the Bacteroidota is independent of the Lol system that traffics lipoproteins to the periplasmic face of the OM^28,50–52^ and that Bacteroidota SLPs use a targeting signal that is distinct from that for the Lol pathway^53,54^.

It is notable that the Bacteroidota LPS translocon achieves SLP transport using a highly canonical LptADE core. This is in marked contrast to the previously-discovered SLP-translocating LptD homologue of *B. burgdorferi* which has diverged extensively from the canonical LptD protein and which is predicted to form a circular 32-stranded barrel with no LptE plug^29^. Freed from the requirement to transport LPS, the *B. burgdorferi* LptD homologue appears to have evolved to provide a large transmembrane pore specialised for SLP export. By contrast, the Bacteroidota LPS translocon has retained the canonical core architecture required for LPS transport while acquiring additional structural elements that facilitate the translocation of complex SLPs.

The LptADE core at the heart of the Bacteroidota LPS translocon is highly conserved across LPS-containing bacteria^6,55^. This raising the possibility that the ability to export SLPs is not unique to the Bacteroidota translocon but is a more general property of LPS translocons. In this model canonical LPS translocons have an intrinsic capacity to translocate small SLPs alongside LPS, while systems such as that of the Bacteroidota have acquired additional components that expand the size and complexity of the SLP substrates they can accommodate. If so, LPS translocons could provide a universal mechanism for SLP trafficking in LPS-containing Gram-negative bacteria.

## Data availability

Cryo-EM density maps and atomic coordinates are deposited in the Electron Microscopy DataBank (EMDB) with the following accession numbers: EMD-79007, EMD-79000, EMD-79001, and EMD-79006 for LPT_Fj_ and EMD-79056, EMD-79055, EMD-79054, and EMD-79053 for LPT_Bt_. Atomic coordinates are deposited in the Protein Data Bank (PDB) with the following accession numbers: PDB 38OL for LPT_Fj_ and PDB 38QP for LPT_Bt_.

SIMPLE4 can be accessed at https://github.com/hael/SIMPLE.

Uncropped gels and immunoblots are provided with this paper.

Requests for materials should be addressed to BCB.

## Acknowledgments

We thank Vaishnavi Ravikumar and Marjorie Fournier of The Advanced Proteomics Facility, Department of Biochemistry, University of Oxford for assistance in proteomics experiments and proteomics data processing, Emilie Alard for help with proteomics sample preparation and data processing, and Raman Dhaliwal and Charlotte Melia for carrying out the TEM analysis. We thank Ian Lidbury and Laila Moushtaq-Kheradmandi for sharing advice regarding *F. johnsoniae* SUS systems. We acknowledge the use of the Advanced Proteomics Facility and the Sir William Dunn School of Pathology Electron Microscopy Facility. This work was supported by European Research Council Advanced Awards 101201241 and 833713, Wellcome Discovery Award 317933/Z/24/Z, and by St. Jude Children’s Research Hospital.

## Author Contributions

XL and LOT carried out all genetic and biochemical experiments. Electron microscopy data were collected by OJB and SML. JC and HE optimised the cryoEM processing workflow. SML determined all structures. XL, LOT, SML, and BCB conceived the project. FKM supervised the proteomics analysis. BCB and SML supervised the project and secured funding. All authors interpreted data and wrote the manuscript.

## Methods

### Bacterial strains and growth conditions

All strains and plasmids used in this work are listed in Supplementary Tables S2 and S3. *F. johnsoniae* was routinely cultured aerobically in Casitone Yeast Extract (CYE) medium^56^ at 30 °C with shaking. For experiments testing growth on complex sugars, cells were cultured in a 96-well plate in a SPECTROstar Nano plate reader using modified minimal A medium^48^ containing 0.25 % (w/v) of either Carob galactomannan (Megazyme, CAS Number: 11078-30-1) or tamarind xyloglucan (Megazyme, CAS Number: 37294-28-3) or glucose as the sole carbon source. Where required the growth medium contained 100 µg ml^-1^ erythromycin, or either 0.2 µg ml^-1^ (liquid culture) or 2 µg ml (agar plates) aTC (CAY10009542, Cambridge Bioscience Ltd).

*B. thetaiotaomicron* was grown anaerobically (10 % CO_2_, 10 % H_2_, 80 % N_2_) at 37 °C in a Don Whitley Scientific A45 workstation. *B. thetaiotaomicron* was cultured in enriched brain–heart infusion medium (EBHI) containing 37 g l^−1^ brain–heart infusion powder (Oxoid, CM1135B), 5 g l^−1^ yeast extract, 1 μg ml^−1^ haemin, 0.5 g l^−1^ cysteine and 1 μg ml^−1^ vitamin K.

*E. coli* strains were grown aerobically in LB medium at 37 °C with shaking, or on LB agar plates. Where appropriate the growth medium contained 100 µg ml^-1^ ampicillin or 50 µg ml^-1^ kanamycin.

### Genetic constructs

Plasmids were constructed by Gibson cloning^57^ using the primers and target DNA in Supplementary Table S4.

Suicide and expression plasmids were introduced into the appropriate *F. johnsoniae* background strain by triparental mating as previously described^58^. Chromosomal modifications were introduced using the suicide vector pYT313 harboring the counter-selectable *sacB* gene as previously described^59^. Strains to enable the depletion of the essential LPT*_Fj_* subunits were constructed as previously described^21^. This method involves placing a copy of the target gene under the control of an aTC-inducible promoter at a phenotypically neutral site in the chromosome then deleting the native copy of the target gene in the presence of aTC to maintain expression of the introduced copy of the gene.

*B. thetaiotaomicron* chromosomal modifications were made in the thymidine kinase-deficient strain VPI-5482 Δ*tdk* using the suicide plasmid pExchange-tdk^60^. Transconjugants were selected on EBHI plates containing 25 μg ml^−1^ erythromycin and 200 μg ml^−1^ gentamicin. Colonies were restreaked on fresh plates of the same medium and resulting single colonies cultured overnight in EBHI without antibiotic selection. Cells that had lost the integrated plasmid were then selected on EBHI containing 200 μg ml^−1^ 5-fluorodeoxyuridine (Thermo Scientific, L16497 ME). Colonies were then restreaked on fresh plates of the same medium and colonies containing the desired mutation identified by PCR.

All plasmid constructs and chromosomal modifications were confirmed by sequencing.

### Immunoblotting

Immunoblotting was carried out as previously described^61^. Antibodies against LPT*_Fj_* subunits, were raised in rabbits against His-tagged recombinant proteins produced using the plasmids listed in Supplementary Table S3. Antisera against BAM_Fj_ subunits, SusCDE subunits, and SkpA were as previously described^21^. The following commercial antisera were used: anti-Strep-tag (34850 Qiagen), anti-GroEL (G6532 Merck), anti-ALFA-Tag (N1582 Synaptic Systems GmbH), anti-His-tag (H1029-100UL Merck Life Science UK Limited), anti-HA-tag (26183 Thermo Fisher Scientific), anti-FLAG-tag (F1804 Merck), anti-mouse IgG peroxidase conjugate (A4416 Merck), and anti-rabbit IgG peroxidase conjugate (31462 Pierce). Antibodies were used at the following dilutions: anti-LptE, anti-LptR, and anti-LptS all 1:100; anti-LptP, anti-His-tag, and anti-FLAG-tag1:1000; anti-BamH, anti-SusC, anti-SusD, anti-SusE, anti-SkpA, anti-Strep-tag, anti-ALFA-tag, anti-HA-tag all 1:3000; anti-GroEL 1:50000.

### LPT_Fj_ subunit depletion experiments

The desired depletion strain was grown overnight in CYE medium supplemented with 0.2 µg ml^-1^ aTC. Cells from 1 ml of the overnight culture were collected, washed once in 1ml CYE, and resuspended in 1 ml of CYE medium. Cells from this sample were then used to inoculate 15 ml of CYE medium, either with or without 0.2 µg ml^-1^ aTC, to OD_600_=0.02. The cells were then cultured aerobically at 30 °C and cell samples periodically collected into SDS sample buffer for subsequent analysis by immunoblotting. Samples for transmission electron microscopy were collected, processed, and imaged as previously described^21^.

### Determination of cell surface exposure of SLPs by protease protection

The strain for analysis was transformed with a plasmid expressing the appropriate epitope-tagged SLP (pXL184 for SusE-his_6_, pXL393 for SusG-FLAG-his_6_, and pXL409 for PorO-FLAG-his_6_). The cells were then grown overnight in CYE supplemented with erythromycin, and for the LptD or LptN depletion strains also with 0.2 µg ml^-1^ aTC. Cells were collected, resuspended in CYE medium, and then used to inoculate 10 ml of erythromycin-containing CYE medium to OD_600_=0.02, supplementing with 0.2 µg ml^-1^ aTC as required. The cells were collected at the desired time point by centrifugation and resuspended to OD_600_=8 in phosphate buffered saline (PBS) containing 10 mM MgCl_2_. A 80 ul sample was supplemented as appropriate with 200 μg ml^−1^ proteinase K (ThermoFisher) and 1% (v/v) Triton X-100 (Merck) and incubated for 20 min at room temperature. Reactions were stopped by the addition of 5 mM phenylmethylsulphonyl fluoride (ITW Reagents) followed by incubation at 100 °C for 5 min, addition of SDS-PAGE sample buffer, and further incubation at 100 °C for 5 min before analysis by immunoblotting.

For the depletion strains the presence of the SLP-producing plasmid and of erythromycin in the growth medium slightly retarded growth relative to the plasmid-free parental strain. Consequently, for the LptN depletion strain the time window in which LptN has become heavily depleted but the cell envelope is intact and LptD levels not appreciably affected is delayed relative to the plasmid-free control. For this strain the proteinase K experiments were carried out on cells taken 5 h after aTC removal, this being phenotypically equivalent to the 4 h time point in the plasmid-free cells. Due to this change, we provide confirmation of the state of the harvested cells in Fig. 5g.

### Purification of LPT_Fj_, LPT_Bt_, BAM_Fj_, and SusCDE complexes

To purify LPT_Fj_ complexes containing Twin-Strep tagged LptD or Twin-Strep tagged LptN the relevant strain was cultured for 22 h in CYE medium using 1 l culture volume in 2.5 l flasks. A total culture volume of 12 l was used for sample preparations for structure determination, and 4 l of culture was used for analytical purifications of LPT*_Fj_* variants. Cells were harvested by centrifugation at 12,000*g* for 30 min and stored at −20 °C until further use. All purification steps were carried out at 4 °C. Cell pellets were resuspended in buffer W (100 mM Tris-HCl pH 8.0, 150 mM NaCl, 1 mM EDTA) containing 30 μg/ml DNase I, 400 μg/ml lysozyme and 1 mM phenylmethylsulfonyl fluoride (PMSF) at a ratio of 5 ml of buffer to 1 g of cell pellet. Cells were incubated on ice for 30 min with constant stirring before being lysed by two passages through a TS series 1.1 kW cell disruptor (Constant Systems Ltd) at 30,000 PSI. Unbroken cells were removed by centrifugation at 20,000*g* for 20 min. The supernatant was recovered and total membranes were collected by centrifugation at 230,000*g* for 75 min. Membranes were resuspended in buffer W to a protein concentration of 6.5 mg/ml and solubilized by incubation with 1% (w/v) lauryl maltose neopentyl glycol (LMNG, Anatrace) for 2 h. Insoluble material was removed by centrifugation at 230,000*g* for 75 min. Endogenous biotin-containing proteins were masked by addition of 1 ml BioLock solution (IBA Lifesciences) per 100 ml of supernatant and incubation for 20 min with constant stirring. The entire solution was then incubated overnight with 1 ml of Strep-TactinXT 4Flow High Capacity resin (IBA Lifesciences) that has had been equilibrated in buffer W containing 0.01% LMNG (buffer WD). The resin was collected into a gravity column and washed with 10 column volumes (CV) of buffer W containing 0.01% LMNG (buffer WD). Bound proteins were eluted with 6 CV Strep-TactinXT BXT buffer (IBA Lifesciences) containing 0.005% LMNG. The eluate was concentrated to 500 μl using a 100-kDa molecular weight cutoff (MWCO) Amicon ultra-15 centrifugal filter unit (Merck) and then injected onto a Superose 6 Increase 10/300 GL column (Cytiva) previously equilibrated in buffer WD. Peak fractions were collected and concentrated using a 100-kDa MWCO Vivaspin 500 column (Sartorius).

Purification of Twin-Strep tagged SusC complexes was carried out by the same protocol from 1 l of cell culture.

To purify LPT_Bt_, 6 ml of an overnight culture of the Twin-Strep tagged LptD strain was used to inoculate fifteen 1 l bottles of pre-warmed EBHI medium. The bottles were then cultured for 7 h to reach an OD_600_∼1.8-2.0. Cells were collected by centrifugation at 6,000 xg for 30 min at 4 °C and stored at −20 °C until further use. LPT_Bt_ was purified from the cell pellets using the same methodology used to isolate LPT_Fj_.

### Peptide mass fingerprinting

Single gel bands samples were excised from Coomassie-stained gels. For whole sample proteomic analysis, SDS-PAGE was carried out only until the sample had fully entered the gel and the protein smear at the top of the gel was excised. Samples were subject to in-gel trypsin digestion and electrospray mass spectrometry at the Advanced Proteomics Facility (University of Oxford, UK).

### Cryo-EM sample preparation and imaging

Purified complexes (3.5 μl each) of *F. johnsoniae* Twin-Strep-LptD or Twin-Strep-LptN complexes (A_280nm_ = 1.0 –1.34) or *B. thetaiotaomicron* Twin-Strep-LptD (A_280nm_ = 1.2) were adsorbed to glow discharged holey carbon-coated grids (Quantifoil 300 mesh, Au R1.2/1.3 for 15 s at 15 mA). Grids were then blotted for 3 s at 100% humidity at 4 °C and frozen in liquid ethane using a Vitrobot Mark IV (FEI). Data were collected on three microscopes: a Titan Krios G3i fitted with a BioQuantum K3 imaging filter and K3 camera operating at 300 kV and at 130,000× magnification, a Krios G4 fitted with a Selectris X imaging filter and a Falcon 4i camera operating at 300 kV and at 130,000× magnification, and an Arctica fitted with a BioQuantum K3 imaging filter and a K3 camera operating at 200 kV and at 130,000× magnification. Pixel sizes ranged from 0.630 Å to 0.724 Å, and movies were collected at a total dose ranging from 45 e^−^ A^−2^ to 54 e^−^ A^−2^ (Extended Data Table 1).

### Cryo-EM data processing

Patched motion correction, contrast transfer function (CTF) parameter estimation, particle picking, extraction and 2D classification were performed in SIMPLE 4.0 (https://github.com/hael/SIMPLE and ^62^). 2D classification used a staged approach with automated particle pruning. All downstream processing was carried out in cryoSPARC (https://doi.org/10.1038/s41592-020-00990-8) or RELION^63^. Global resolution was estimated from gold-standard Fourier shell correlations using the 0.143 criterion, and local resolution estimation was calculated within cryoSPARC using a Fourier shell correlation threshold of 0.5.

The cryo-EM processing workflow for the *F. johnsoniae* LPS translocon complex is outlined in Extended Data Fig. 1. In brief, particles were subjected to a second round of reference free 2D classification in cryoSPARC. Three volumes were then generated from a 554,003 particle subset of the 2D-cleaned particles by multi-class ab initio reconstruction using a maximum resolution cut-off of 12 Å. Particles (224,707) from the most populated and structurally well-defined class were selected and non-uniform refined against its corresponding volume, yielding a 3.24 Å map. Bayesian polishing was performed in RELION followed by local and global CTF refinement (fitting beamtilt and trefoil) and a further round of non-uniform refinement in cryoSPARC yielding a 2.93 Å map. Local refinements were subsequently performed using masks focused on either the membrane and extracellular region or the periplasmic region. Focused 3D classification using these masks yielded volumes with improved densities for extramembrane and periplasmic components, respectively. A second local refinement was performed on each volume using their respective focus mask, producing maps at 3.10 Å and 3.47 Å resolution. Maps were sharpened using deepEMhancer (https://doi.org/10.1038/s42003-021-02399-1) and the resulting reconstructions were combined into a composite map for model building and refinement.

The cryo-EM processing workflow for the *B. thetaiotaomicron* LPS translocon complex is outlined in Extended Data Fig. 3. In brief, particles were subjected to a second round of reference free 2D classification in cryoSPARC. Five volumes were then generated from a 967,983 particle subset of the 2D-cleaned particles by multi-class ab initio reconstruction using a maximum resolution cut-off of 12 Å. Particles (217,663) from the most populated and structurally well-defined class were selected and non-uniform refined against its corresponding volume, yielding a 3.47 Å map. Bayesian polishing was performed in RELION followed by local and global CTF refinement (fitting beamtilt and trefoil) and a further round of non-uniform refinement in cryoSPARC yielding a 3.27 Å map. Local refinements were subsequently performed using masks focused on either the membrane and extracellular region or the periplasmic region. Focused 3D classification using these masks yielded volumes with improved densities for extramembrane and periplasmic components, respectively. A second local refinement was performed on each volume using their respective focus mask, producing maps at 2.94-Å and 3.47-Å resolution. Maps were sharpened using deepEMhancer and the resulting reconstructions were combined into a composite map for model building and refinement.

### Biotin labelling of cell surface proteins

Overnight cultures of the strains to be analysed were inoculated into CYE medium at a starting OD_600_=0.02 and then cultured to OD_600_=0.5. 12 ml samples of the cultures were collected by centrifugation at 8,000*g* at 4 °C for 10 min. The cell pellets were washed twice with ice cold PBS, resuspended with 1 ml cold PBS to OD_600_=6, and then labelled with 0.2 mM EZ-LinkTM sulfo NHS-LC-LC-Biotin (21338, Thermo Fisher) for 2 min at room temperature. The labelling reaction was terminated by the addition of 1M Tris-HCl pH7.5 to a final concentration of 100 mM, and then incubating for 10 min at room temperature. Cells were collected by centrifugation at 8,000*g* at 4 °C for 5 min and then washed three times with cold PBS. Cell pellets were stored at −20 °C at this page.

The frozen cell pellets were resuspended with 1 ml PBS supplemented with 1% SDS and then incubated at 100 °C for 10 min. The samples were then centrifuged at 20,000*g* for 20 min at room temperature. The resulting supernatant fractions were collected, diluted with PBS buffer to give a final SDS concentration of 0.1%, and then mixed with 30 μl high capacity Neutravidin agarose resin (29202, Thermo Scientific) that had been pre-equilibrated with PBS buffer containing 0.1% SDS. The samples were then incubated for 1 h at room temperature on a roller. After this, the resin was recovered by centrifugation at 200*g* for 3 min, washed 3 times with 3 ml PBS buffer containing 0.1% SDS, resuspended in 100 μl PBS buffer containing 4% SDS, and incubated at 100 °C for 10 min. The resin was removed by centrifugation and the resulting supernatant retained for analysis by MS proteomics, SDS-PAGE and immunoblotting as required.

### Preparation of biotinylated samples for proteomics MS

Samples were prepared using a filter aided sample preparation (FASP) workflow^64^, with some modifications. Briefly, the SDS-solubilized samples were reduced with 10 mmol l^-1^ dithiothreitol (DTT) for 30 min at room temperature (RT), followed by alkylation with 55 mmol l^-1^ iodoacetamide (IAA) for 30 min at room temperature protected from light. Samples were then centrifuged at 14,000*g* in a bench top centrifuge for 3 min to remove insoluble material.

Vivacon 500, 30kDa Hydrosart filter units (Sartorius) were pre-equilibrated with 450 µl of UA buffer (8 M urea, 100 mM Tris-HCl pH 8.8) and centrifuged at 14,000*g* for 30 s. Filters showing substantial buffer passage were discarded. Up to 60 µl of reduced and alkylated sample supernatant was loaded onto the UA buffer on the filter tops and centrifuged at 14,000*g* for 15 min. The flow-through was discarded and the filters were subsequently washed three times with 200 µl of UA buffer, followed by three times with 200 µl of 100 mM triethylammonium bicarbonate (TEAB) buffer (90114, Thermo Fisher) buffer, through successive rounds of buffer addition and centrifugation at 14,000*g* for 10 min, discarding the flow-through.

Filters were then transferred to clean collection tubes, and proteins were digested with trypsin (MS grade, Thermo Fisher) diluted in 50 µl of freshly prepared 40 mmol l^-1^ ammonium bicarbonate (ABC) buffer at a 1:100 enzyme-to-protein (w/w) ratio. Samples were mixed at 600 rpm for 1 min and incubated at 37 °C overnight under humidified conditions.

Peptides were recovered by centrifugation at 14,000*g* for 10 min and collection of the flow-through. Filters were then washed twice with 50 µl ABC buffer and once with 50 µl 0.5 mol l^-1^ NaCl, followed by centrifugation at 14,000*g* for 10 min after each wash. All flow-through fractions were collected and combined. Finally, samples were acidified by the addition of 1% trifluoroacetic acid (TFA). 4% acetonitrile (AcN) was added to the peptide mix at a 1:1 (v/v) ratio.

Peptide samples were loaded onto Evotip Pure tips (Evosep Biosystems) according to the manufacturer’s instructions. Briefly, Evotips were rinsed with 20 µl solvent B (0.1% formic acid (FA) in AcN), centrifuged at 800*g* for 1 min, pre-wetted with 1-propanol, and equilibrated with 20 µl solvent A (0.1% FA in water), followed by centrifugation at 800*g* for 1 min. Peptide samples were loaded onto the equilibrated Evotips and centrifuged at 800*g* for 1 min. Evotips were subsequently washed with 20 µL solvent A and centrifuged at 800*g* for 1 min. Finally, 100 µL solvent A was added and the tips were centrifuged at 800*g* for 10 s to maintain hydration prior to LC–MS/MS analysis.

### Preparation of whole cell samples for proteomics analysis

For whole cell proteomic analysis of LPT variants 1 ml of exponentially growing cultures (OD_600_=0.5) were collected by centrifugation at 8,000*g*, and cell pellets were washed with ice cold PBS and stored at −20 °C. Cell pellets were resuspended in lysis buffer containing 1 % sodium dodecyl sulfate, 0.1 M ammonium bicarbonate pH 8.0 and cOmplete protease inhibitor cocktail (Roche). Samples were vortexed well and sonicated 5 times in a water bath for 1 min, with 1 min incubation on ice between each pulse cycle. This was followed by the addition of 2 units of DNase (NEB), incubation on ice for 30 min and centrifugation at 4 °C for 30 min at 12,700 rpm. The supernatant collected and sample clean-up (detergent removal) was performed based on the SP3 methodology^65^. Briefly, samples were reduced for 30 min at room temperature using 10 mM TCEP (tris(2-carboxyethyl)phosphine), followed by alkylation for 30 min in the dark at room temperature using 2-chloroacetamide. Subsequently, SpeedBeads Magnetic Carboxylate Modified Particles (GE Healthcare) that had been washed 3 times in HPLC-grade water, were mixed with the sample in a 10:1 ratio (bead:sample). Binding was aided using ≥ 50 % ethanol. Samples were incubated for 15 min at 1000 rpm at 24 °C. Beads were then washed 5 times with 80 % ethanol. Digestion buffer (100 mM ammonium bicarbonate) was added to the washed beads and pre-digestion with endoprotease LysC (Wako; 1:100) was carried out at 37 °C for 2 h at 1000 rpm. This was followed by 18 h digestion with trypsin (Promega, 1:40) at 37 °C at 1000 rpm. The supernatant was acidified (final concentration of 5 % formic acid) and collected post digestion. 200 ng equivalent of digested peptides were further desalted using an Evotip Pure (Evosep) as described above.

### Proteomics analysis

Mass spectrometry analysis of Evotip loaded samples was performed with an Orbitrap Astral ZOOM (Thermo) coupled to an Evosep Eno LC system.

For the biotin-labelling experiments, peptides were resolved on an EV1182 Evosep Performance Column, using a 200 Sample Per Day (200 SPD) method. For the whole cell proteomics analysis, peptides were separated on an EV1109 (Evosep) analytical column using a 60 SPD method The mass spectrometer was operated in positive-ion mode using nanospray ionisation, with the spray voltage set to 1.9 kV and the ion transfer tube temperature maintained at 290 °C. MS was performed in Data Independent Acquisition (DIA) mode(Guzman, Rykaer et al. 2026). DIA acquisition was performed with automatic window placement and a 0.6-s time-controlled acquisition loop. MS1 spectra were recorded in profile mode, whereas DIA MS/MS spectra were recorded in centroid mode. A full-MS resolution of 240,000, a scan range of 380 to 980 m/z, and a full-MS AGC target value of 5 × 10⁶ with a maximum injection time of 5 ms was applied. Fragment ion scans were performed at 80,000 resolution, over an m/z range of 150–2000, with a maximum Injection Time of 3 ms and an AGC target of 5 × 10⁴. A total of 300 windows with equal m/z widths of 2 were used for ion isolations. Window-placement optimization was enabled. The isolated ions were fragmented using higher-energy collisional dissociation (HCD) with 25% normalized collision energy. MS raw files were subsequently searched against an in silico generated tryptic spectral library from *Flavobacterium johnsoniae* UW101 proteome, downloaded from UniProt (2026.02.28), using DIA-NN (versions 2.2.0 or 2.6.0)^66^.Common contaminants in-built within the software’s algorithm were included. Match between runs was enabled. Identification FDR was set at 1 %. All other default parameters were maintained.

All downstream proteomics data analysis was performed in Perseus 2.1.3.0^67^. Briefly, the DIA-NN estimated label-free intensity values were Log2 transformed and normalised by median subtraction. Proteins not identified in all biological replicates were filtered, and the biotin-labelled cell surface proteome values were then subtracted from their corresponding whole cell proteome values to correct for expression differences. Significantly changing proteins between wild-type and mutant strains were identified using a 2-sample 2-tailed t-test analysis with permutation-based FDR cut-off of <0.05 and a S0 of 0.1.

The proteins detected in the MS experiments were categorised as follows. Proteins with signal peptides or lipoprotein signal peptides were first extracted using SignalP 6.0^68^ to obtain datasets containing either outer membrane proteins (OMPs) plus periplasmic proteins, or lipoproteins. Lipoproteins were then manually sorted between SLPs and periplasmic lipoproteins (PLPs). Lipoproteins were classified as SLPs if they were either known SLPs, or were SusD homologues, or if they were coded at a polysaccharide utilisation locus (PUL), of if they contained a lipoprotein export signal (LES)^53^ defined as a net negative charge in the six residues following the lipidated amino terminal cysteine residue. Known PLPs were also excluded. The remaining lipoproteins were assigned as PLPs.

---

**Extended Data Fig. 1:**
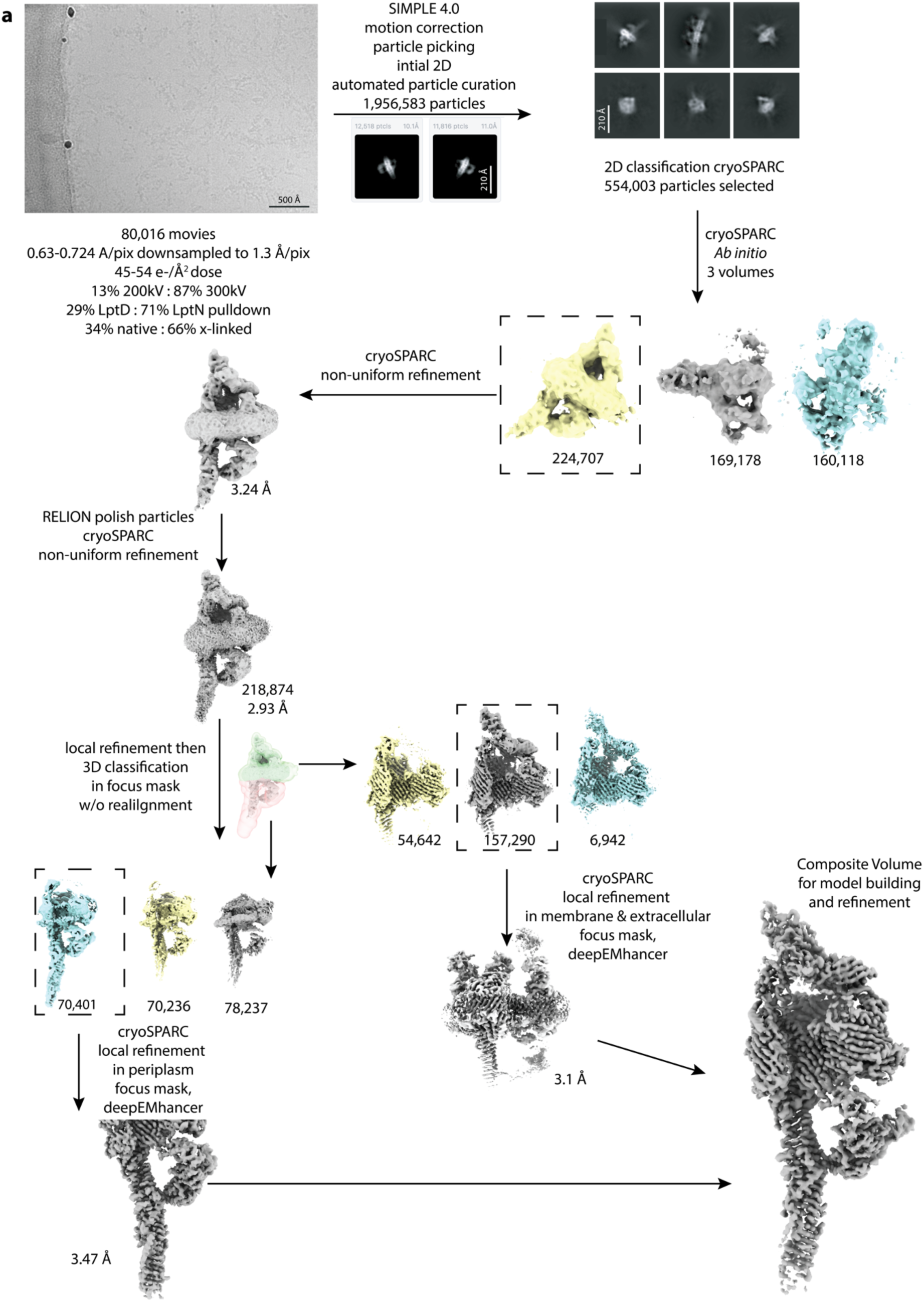
Workflow for the cryoEM analysis of the *F. johnsoniae* LPS translocon complex.

**Extended Data Fig. 2:**
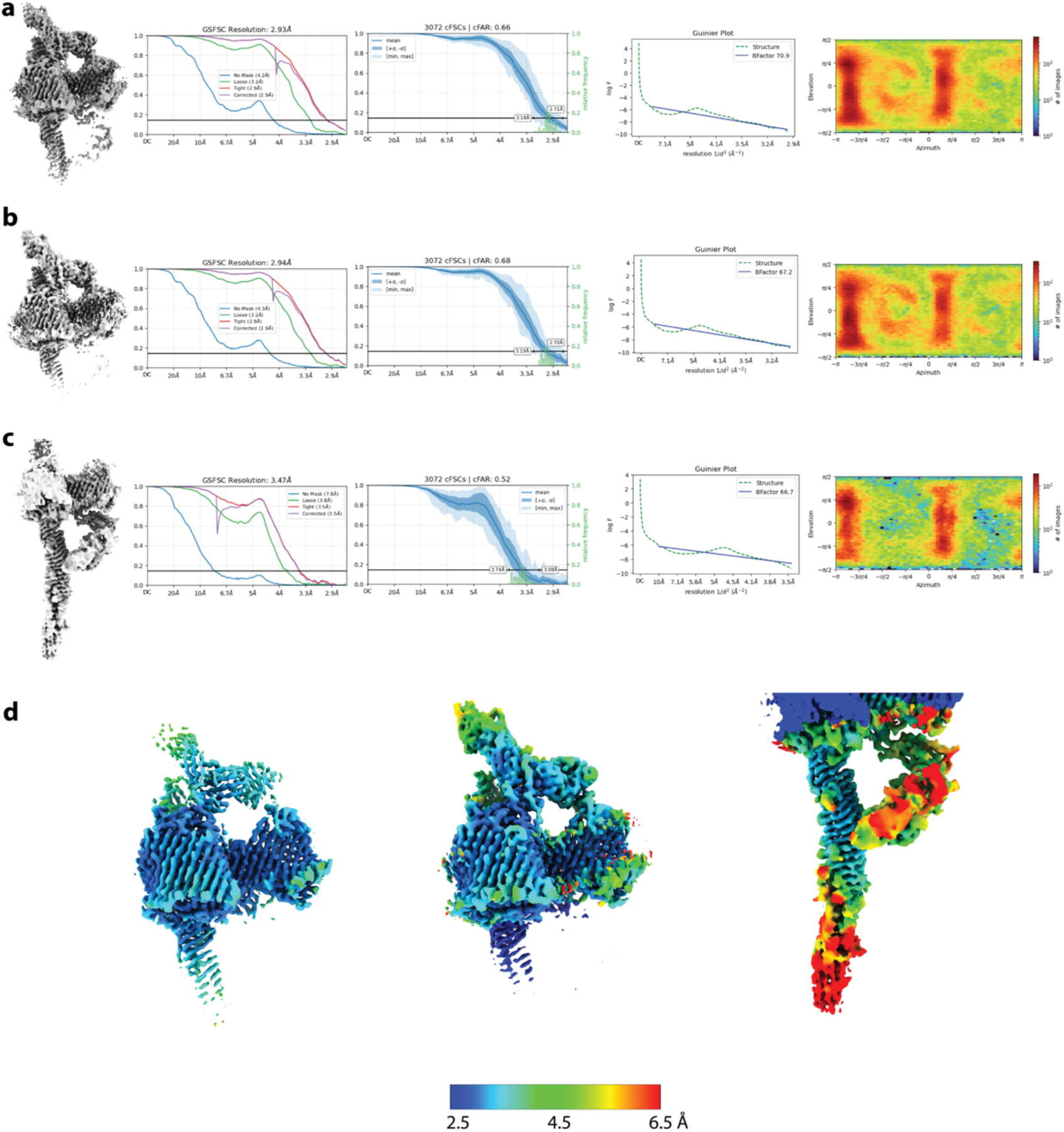
*F. johnsoniae* LPS translocon complex cryoEM volume quality metrics. a-c, From left to right in each panel are the volume / Gold standard Fourier Shell Correlation (FSC) curves used for global resolution estimation / cFSCs (Half-angle: 20^°^), with autotight mask and overlaid frequency plot / Gunier plot with derived B factor used for isotropic sharpening / viewing direction distribution for aligned particles **a,** Consensus volume. **b,** Membrane and extracellular region focus masked. **c,** Periplasmic region focus masked. **d,** Local resolution-coloured volumes for the consensus volume (left), focus region membrane and extracellular (centre), and focus region periplasm (right).

**Extended Data Fig. 3:**
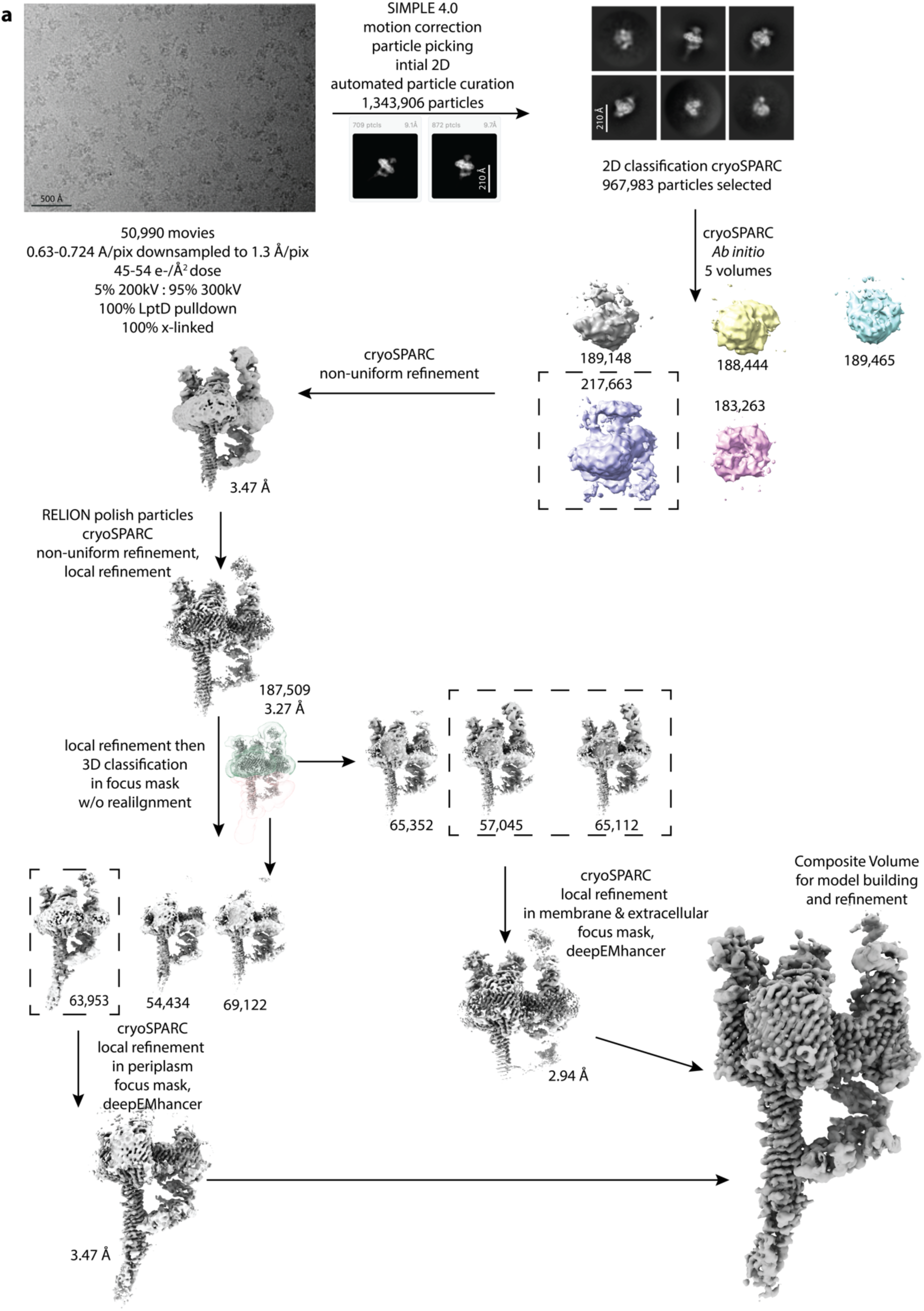
Workflow for the cryoEM analysis of the *B. thetaiotaomicron* LPS translocon complex.

**Extended Data Fig. 4:**
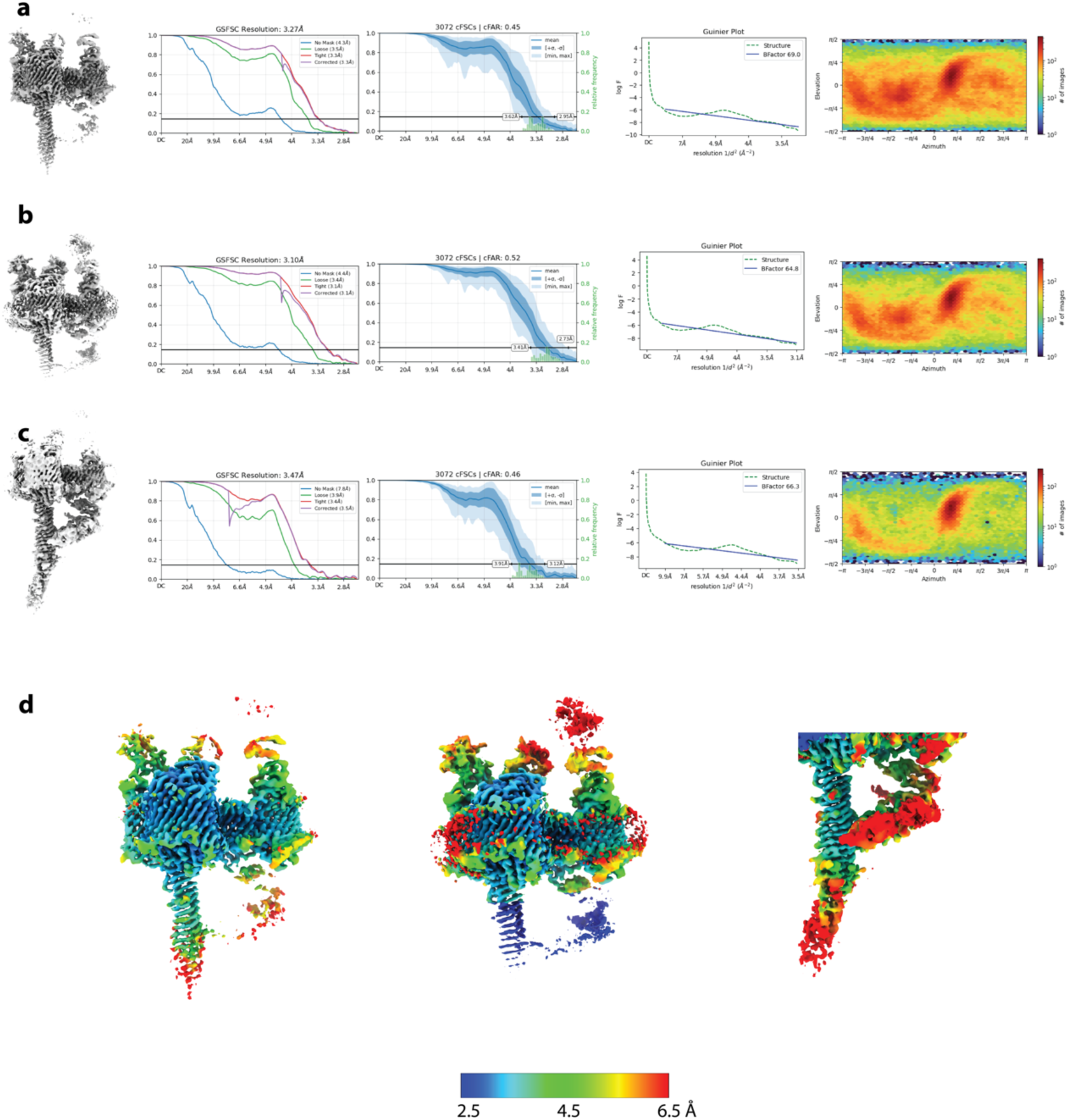
*B. thetaiotaomicron* LPS translocon complex cryoEM volume quality metrics. a-c, From left to right in each panel are the volume / Gold standard Fourier Shell Correlation (FSC) curves used for global resolution estimation / cFSCs (Half-angle: 20^°^), with autotight mask and overlaid frequency plot / Gunier plot with derived B factor used for isotropic sharpening / viewing direction distribution for aligned particles **a,** Consensus volume. **b,** Membrane and extracellular region focus masked. **c,** Periplasmic region focus masked. **d,** Local resolution-coloured volumes for the consensus volume (left), focus region membrane and extracellular (centre), and focus region periplasm (right).

**Extended Data Fig. 5:**
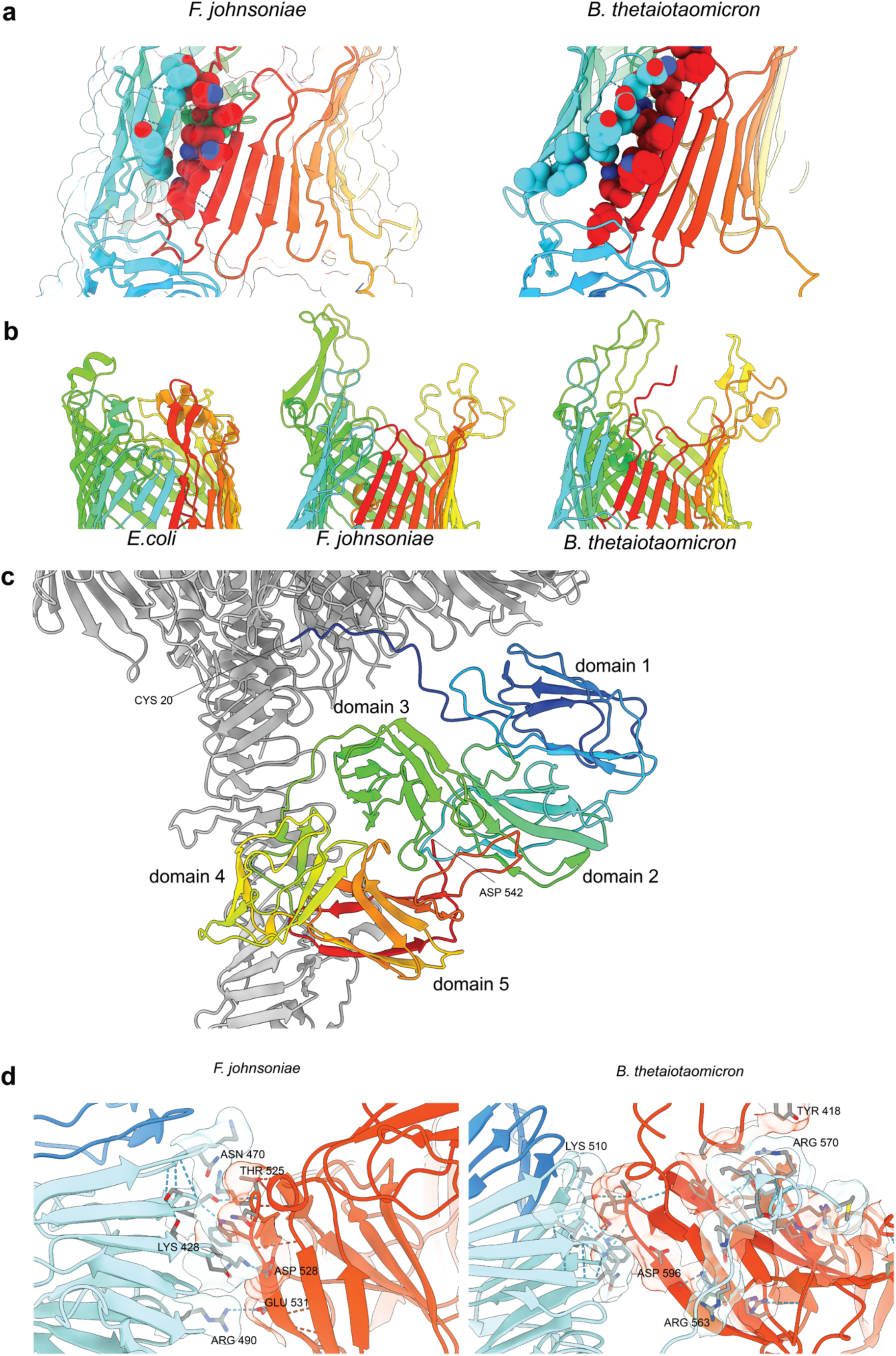
Further structural details of the LptD and LptP subunits. a, The lateral gate of the Bacteroidota LptD barrel is closed by the side chains of the N- and C-terminal strands. A rainbow coloured cartoon of LptD focused on the lateral gate with space-filling representations of side chains that pack across the N and C-terminal strands to close the gate in the absence of hydrogen bonding between the strands. **b,** Comparison of the extracellular loops of the LptD barrels of *E. coli*, *F. johnsoniae*, and *B. thetaiotaomicron* showing that these only clamp down onto LptE in the *E. coli* protein. Proteins are shown in rainbow-coloured cartoon representation. **c,** LptP in the context of the *F. johnsoniae* LPS translocon assembly. LptP is show in cartoon representation and coloured from blue at the N-terminus to red at the C-terminus. **d,** Close-up of the contact site between LptP and LptA in the two Bacteroidota LPS translocons. Different contacts stabilise the interaction between domain 5 of LptP and LptA in the two complexes.

**Extended Data Fig. 6:**
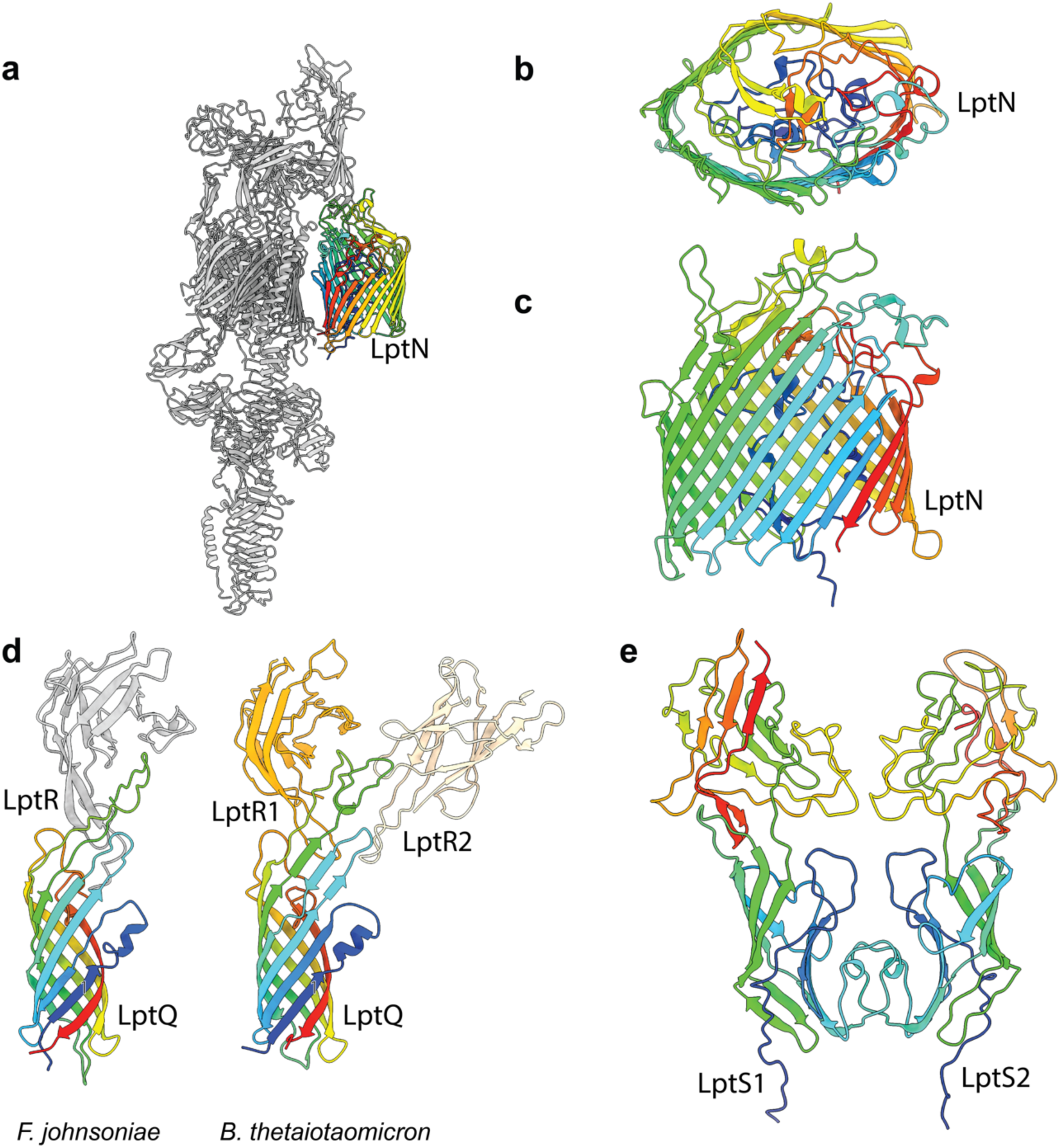
Structural details of the LptN, LptQ, and LptR subunits. a-c, Rainbow coloured cartoon of *F. johnsoniae* LptN. **a,** In the context of the whole LPS translocon complex. **b,** Viewed from the cell exterior. **c,** Viewed from within the plane of the membrane. **d,** Cartoon representation of the LptQR sub-complexes in the Bacteroidota LPS translocons viewed from within the plane of the membrane. LptQ is rainbow coloured. **e,** Structure of the LptS1-LptS2 dimer within the *F. johnsoniae* LPS translocon complex shown in rainbows cartoon representation and viewed from the periplasmic side.

**Extended Data Fig. 7:**
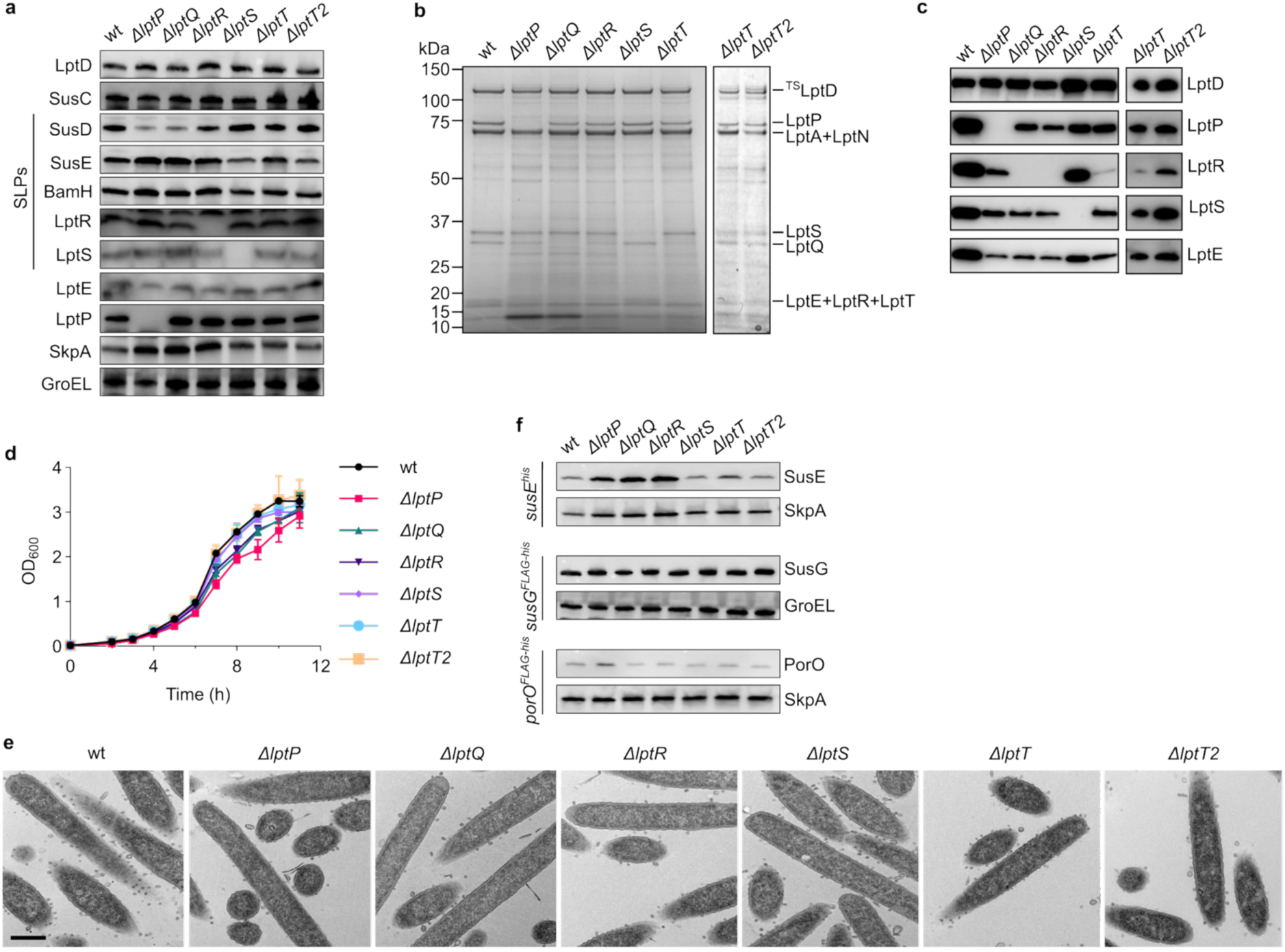
Phenotypic effects of removing non-essential LPT_Fj_ subunits. wt, wild type. **a,** Whole cell immunoblots. SLPs, surface lipoproteins. SkpA is a periplasmic protein. GroEL is a cytoplasmic protein. Similar data were obtained for three biological repeats. **b,c** Isolation of LptD complexes from backgrounds missing the indicated Lpt subunits. Strep-tagged LptD complexes were purified by Streptactin affinity chromatography and analysed by (**b**) Coomassie-stained SDS–PAGE gel and (**c**) Immunoblotting with the available subunit-specific antibodies. Similar results were obtained from 2 biological repeats. **d,** Growth curves on rich CYE medium. Shown are the means ± 1 SD from three biological repeats. **e,** Representative transmission electron microscopy images. Scale bar, 500 nm. **f,** Whole cell immunoblots of the cells used in Fig. 5d showing no reduction in SusE, SusG or PorO levels in the mutants. SkpA, GroEL are loading controls.

**Extended Data Fig. 8:**
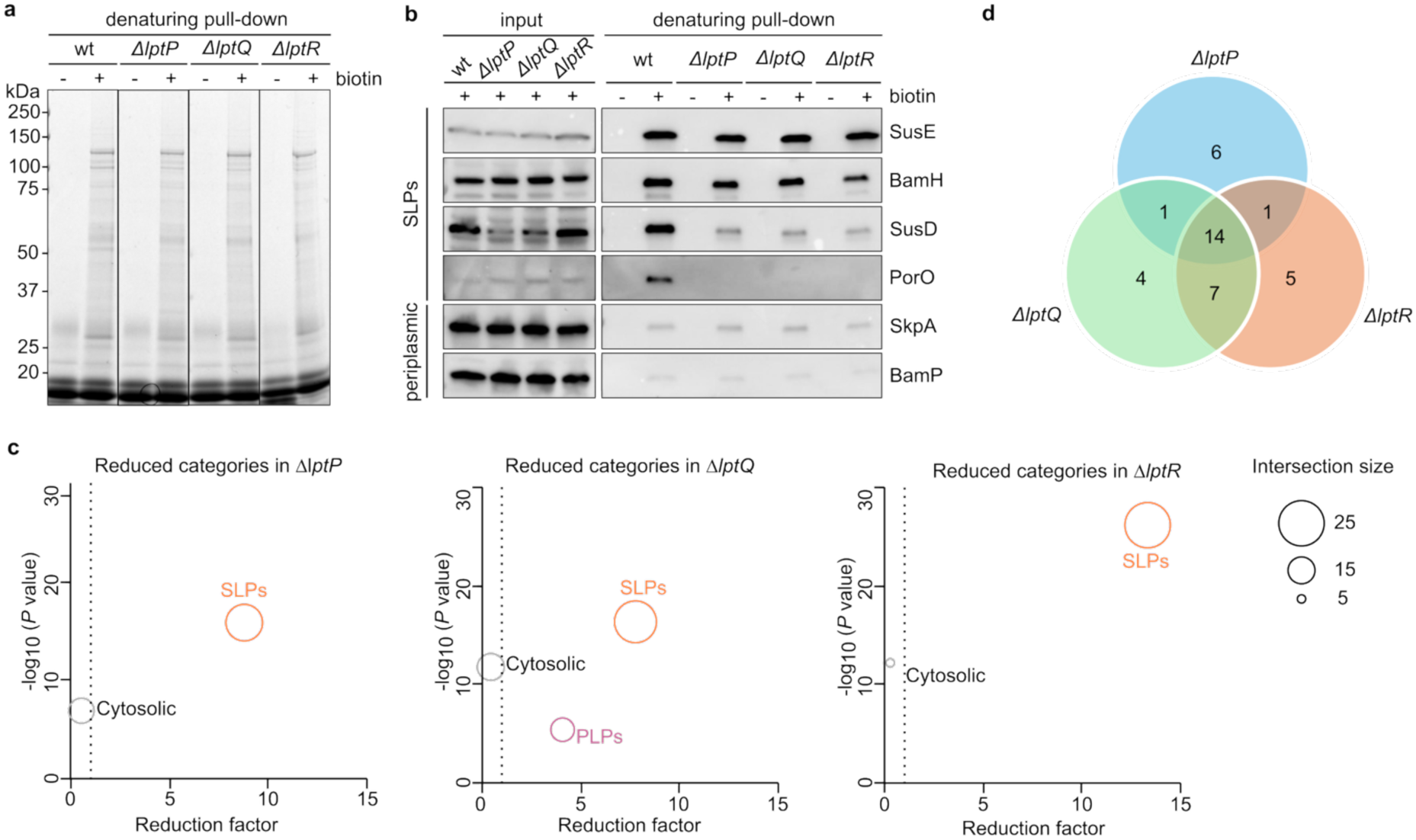
Effects of LPT_Fj_ subunit deletions on the surface exposure of the *F. johnsoniae* SLP proteome. **a,b**, Validation of the NHS-biotin labelling methodology. Strains transformed with a plasmid expressing epitope-labelled PorO were labelled where indicated with sulfo-NHS-LC-LC-biotin (‘biotin’). Biotin-conjugated proteins were then purified under denaturing conditions and analysed by (**a**) Coomassie blue-stained SDS-PAGE and (**b**) immunoblotting with the indicated antibodies or with anti-FLAG in the case of PorO. Similar results were obtained from two biological repeats. wt, wild type. **c,d,** Additional analysis of the datasets characterized in Fig. 5e. **c,** Category enrichment analysis of proteins that show reduced surface exposure in the indicated mutants, using Fisher’s exact test with a Benjamini–Hochberg FDR cut-off < 0.02. Each circle represents an enriched category, with circle size being proportional to the number of enriched proteins. SLPs, surface lipoproteins; PLPs, periplasmic lipoproteins. The dotted line indicates an enrichment factor of 1. **d,** Venn diagram of SLPs that exhibit reduced surface exposure in the indicated mutants at the significance threshold used in Fig. 5e. The identities of the proteins are given in Extended Data Table 3.

**Extended Data Fig. 9:**
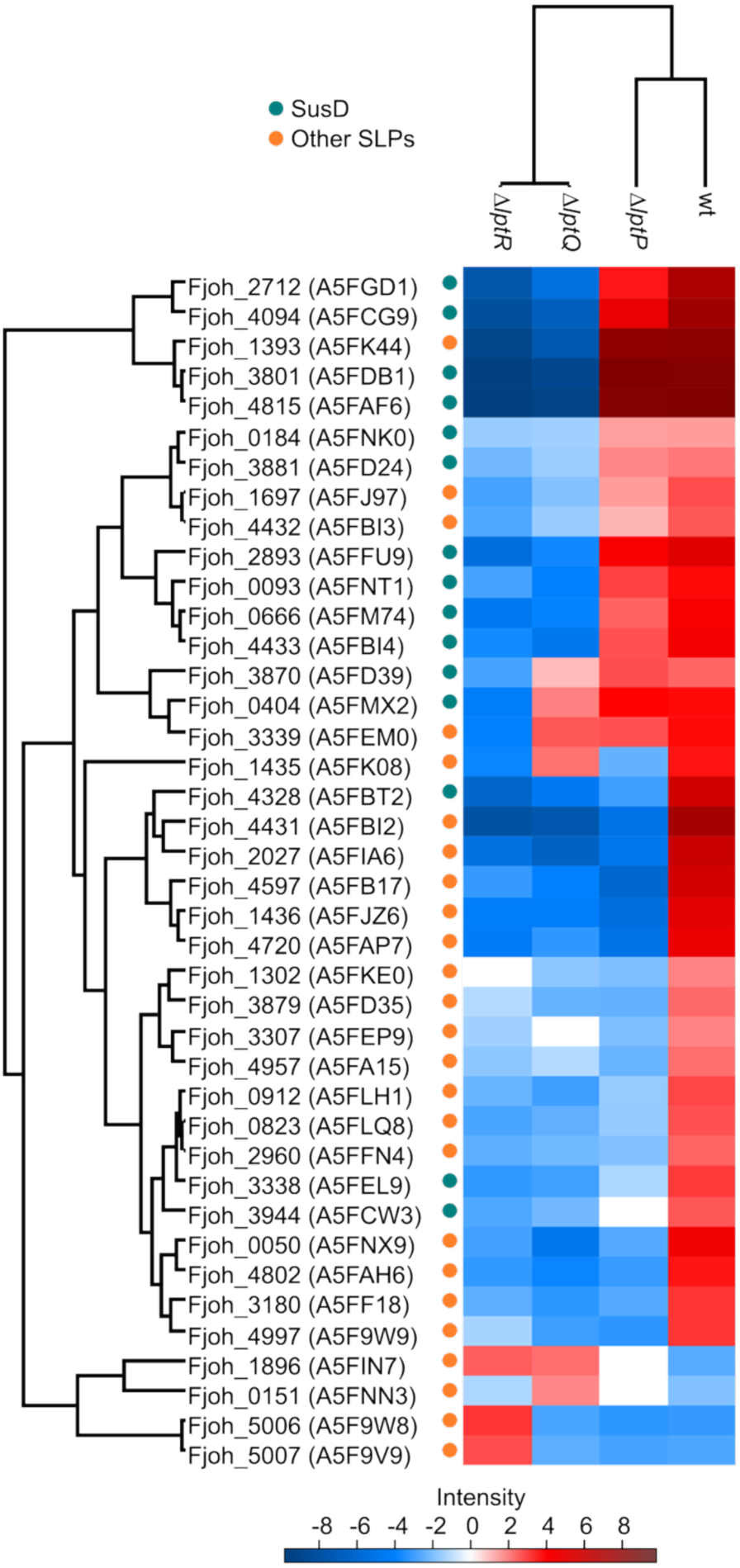
Hierarchical protein clustering analysis of the effects of LPT_Fj_ subunit deletions on the surface exposure of the *F. johnsoniae* SLP proteome. Heat map of change in cell surface exposure of the listed ANOVA-significant SLP proteins across the indicated strains, following ANOVA analysis with post-hoc testing and hierarchical clustering of the entire database. Colours indicate HSD (honestly significant difference) values according to the intensity panel. An additional coloured dot indicates whether proteins are SusD family SLPs or other SLPs.

**Extended Data Table 1:** Cryo-EM data collection, refinement and validation statistics.

|  | *F. johnsoniae*<br>LptD complex<br>(EMDB-79007,79000,79001,79006)<br>(PDB 38OL) | *B. thetaiotaomicron*<br>LptD complex<br>(EMDB-79056,79055,7905<br>4,79053)<br>(PDB 38QP) |
| --- | --- | --- |
| **Data collection and processing** |  |  |
| Magnification | 130,000 / 130,000 / 165,000 |  |
| Voltage (kV) | 200 / 300 / 300 |  |
| Electron exposure (e<sup>-</sup>/Å<sup>2</sup>) | 45 – 54 |  |
| Defocus range (μm) | -0.6 – -2.1 |  |
| Pixel size (Å) | 0.63 / 0.6485 / 0.724 |  |
| Symmetry imposed | C1 |  |
| Initial particle images (no.) | 1,956,583 | 1,343,906 |
| Final particle images (no.) | 218,874 | 187,509 |
| Focus extracellular/membrane | 157,290 | 130,464 |
| Focus periplasm | 70,401 | 63,953 |
| Map resolution (Å) | 2.93 | 3.3 |
| Focus extracellular/membrane | 2.94 | 3.10 |
| Focus periplasm | 3.47 | 3.47 |
| FSC threshold | 0.143 | 0.143 |
| **Refinement** |  |  |
| Model resolution (Å) | 1.8 | 1.9 |
| FSC threshold | 0.143 | 0.143 |
| CC (mask) | 0.77 | 0.75 |
| Mean CC for ligands | 0.70 | 0.63 |
| Model composition |  |  |
| Non-hydrogen atoms | 29,227 | 30,736 |
| Protein residues | 3,577 | 3,676 |
| Ligands | 16 | 27 |
| *B* factors (Å<sup>2</sup>) |  |  |
| Protein | 95 | 170 |
| Ligand | 120 | 133 |
| R.m.s. deviations |  |  |
| Bond lengths (Å) | 0.005 | 0.003 |
| Bond angles (°) | 0.6 | 0.9 |
| Validation |  |  |
| MolProbity score | 3.25 | 3.3 |
| Clashscore | 47 | 63 |
| Poor rotamers (%) | 3.5 | 5 |
| Ramachandran plot |  |  |
| Favored (%) | 83 | 90 |
| Allowed (%) | 16 | 10 |
| Disallowed (%) | 1 | 0 |

**Extended Data Table 2:**
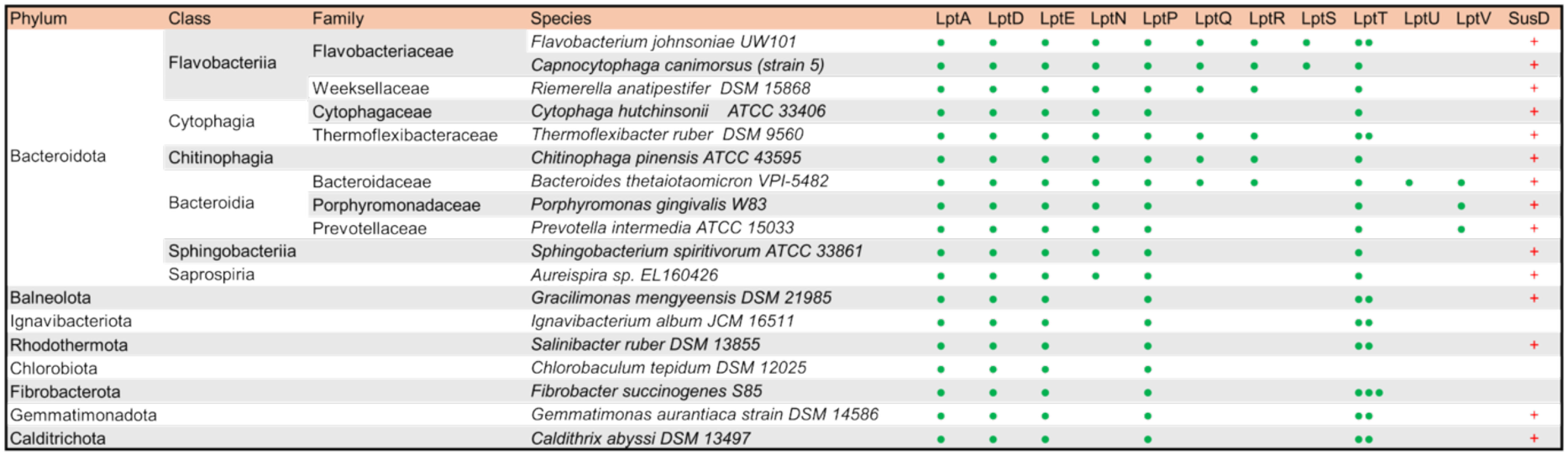
Phylogenetic distribution of *F. johnsoniae* and *B. thetaiotaomicron* LPS translocon subunit homologues. Distribution of LPT*_Fj_* and LPT*_Bt_* subunit homologues within representative members of the Bacteroidota and wider FCB superphylum. Each paralogue is indicated by a green dot. Note that one of the three LptT paralogues in *F. succinogenes* is a tandem fusion of two LptT domains. The presence of members of the SusD protein family (red +) is also shown. See also Supplementary Table 1 for the accession numbers of the proteins identified in this Table.

**Extended Data Table 3:**
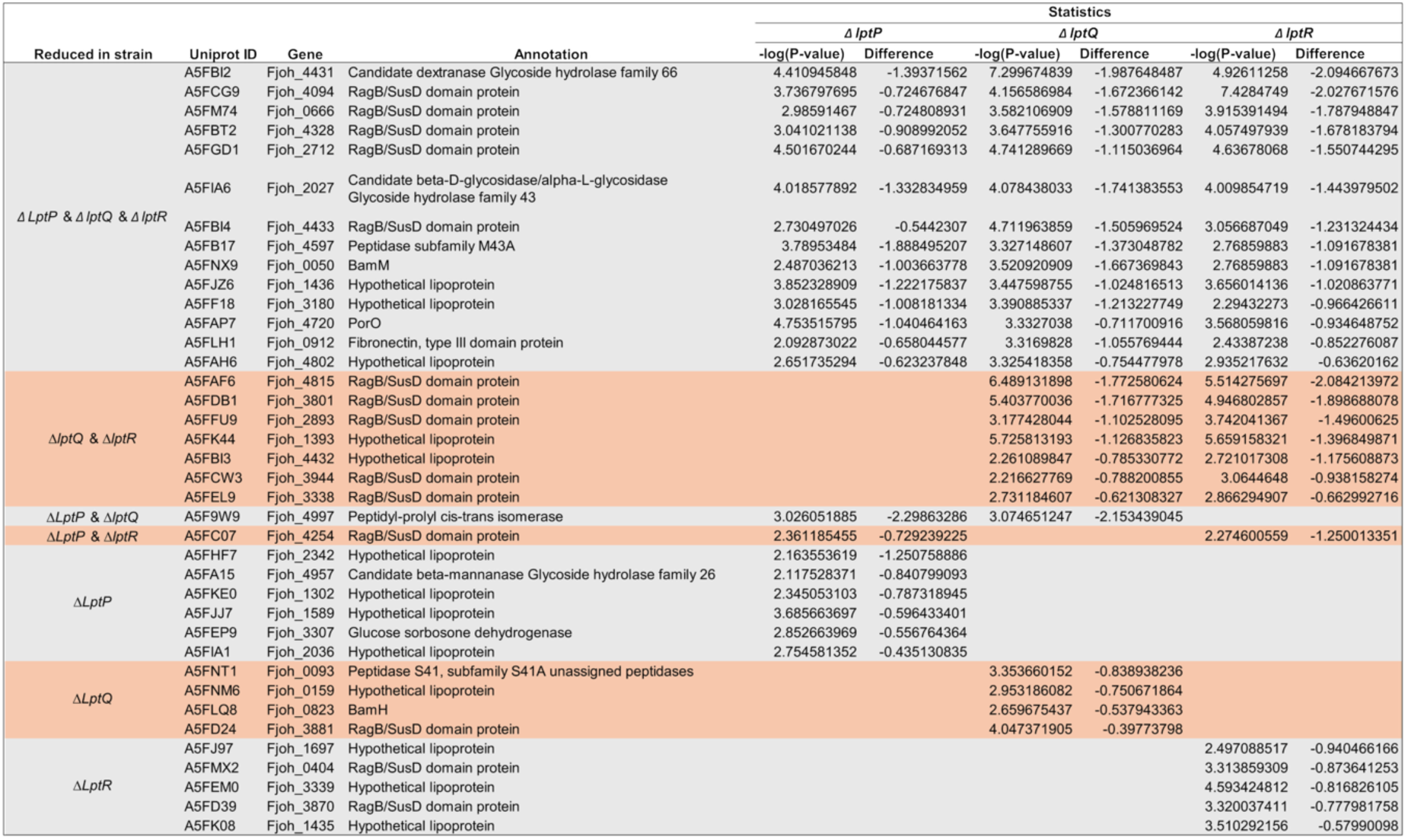
Identities of SLPs that show significantly reduced surface labelling in strains deleted for at least one of LptP, LptQ, or LptR. The significance threshold is that used in Fig. 5e.

