## Supplementary Information for "An expanded LPS translocon mediates surface lipoprotein export in the Bacteroidota"

### Table S1 | Accession numbers of the Lpt subunits identified in Extended Data Table 2.

| Species | LptA | LptD | LptE | LptN | LptP |
|---|---|---|---|---|---|
| *Flavobacterium johnsoniae UW101* | Fjoh_2592 | Fjoh_4586 | Fjoh_1847 | Fjoh_1926 | Fjoh_0663 |
| *Capnocytophaga canimorsus (strain 5)* | Ccan_06900 | Ccan_16770 | Ccan_01410 | Ccan_17620 | Ccan_03570 |
| *Riemerella anatipestifer DSM 15868* | RA0C_1913 | RA0C_1121 | RA0C_1605 | RA0C_1581 | RA0C_1054 |
| *Cytophaga hutchinsonii ATCC 33406* | CHU_3019 | CHU_0749 | CHU_1823 | CHU_1803 | CHU_3504 |
| *Thermoflexibacter ruber DSM 9560* | SAMN04488541_10323 | SAMN04488541_101973 | SAMN04488541_101930 | SAMN04488541_100547 | SAMN04488541_101233 |
| *Chitinophaga pinensis ATCC 43595* | Cpin_6193 | Cpin_6190 | Cpin_5918 | Cpin_6600 | Cpin_0895 |
| *Bacteroides thetaiotaomicron VPI-5482* | BT_3849 | BT_2133 | BT_4376 | BT_0328 | BT_3341 |
| *Porphyromonas gingivalis W83* | PG_0414 | PG_1405 | PG_0147 | PG_0362 | PG_0081 |
| *Prevotella intermedia ATCC 15033* | D2S45_03820 | D2S45_06910 | D2S45_06070 | D2S45_07400 | D2S45_05370 |
| *Sphingobacterium spiritivorum ATCC 33861* | HMPREF0766_10204 | HMPREF0766_10130 | HMPREF0766_10592 | HMPREF0766_11806 | HMPREF0766_10937 |
| *Aureispira sp. EL160426* | AsAng_0020450 | AsAng_0039720 | AsAng_0017630 | AsAng_0046780 | AsAng_0050210 |
| *Gracilimonas mengyeensis DSM 21985* | SAMN06265219_11915 | SAMN06265219_105156 | SAMN06265219_11544 | no | SAMN06265219_11454 |
| *Ignavibacterium album JCM 16511* | IALB_1892 | IALB_1054 | IALB_0614 | no | IALB_1449 |
| *Salinibacter ruber DSM 13855* | SRU_1361 | SRU_0784 | SRU_2651 | no | SRU_0507 |
| *Chlorobaculum tepidum DSM 12025* | CT2096 | CT0523 | CT1770 | no | CT2013 |
| *Fibrobacter succinogenes S85* | Fisuc_1241 | Fisuc_2525 | Fisuc_0376 | no | Fisuc_1637 |
| *Gemmatimonas aurantiaca strain DSM 14586* | GAU_1619 | GAU_1938 | GAU_1569 | no | GAU_1282 |
| *Caldithrix abyssi DSM 13497* | Calab_3123 | Calab_0089 | Calab_3384 | no | Calab_2056 |

| Species | LptQ | LptR | LptS | LptT | LptU | LptV |
|---|---|---|---|---|---|---|
| *Flavobacterium johnsoniae UW101* | Fjoh_2498 | Fjoh_2499 | Fjoh_2510 | Fjoh_0204; Fjoh_1207 | no | no |
| *Capnocytophaga canimorsus (strain 5)* | Ccan_06090 | Ccan_06100 | Ccan_11470 | Ccan_17330 | no | no |
| *Riemerella anatipestifer DSM 15868* | RA0C_1337 | RA0C_1336 | no | RA0C_0331 | no | no |
| *Cytophaga hutchinsonii ATCC 33406* | no | no | no | CHU_2562 | no | no |
| *Thermoflexibacter ruber DSM 9560* | SAMN04488541_102552 | SAMN04488541_102553 | no | SAMN04488541_10027; SAMN04488541_101874 | no | no |
| *Chitinophaga pinensis ATCC 43595* | Cpin_0160 | Cpin_0159 | no | Cpin_2029 | no | no |
| *Bacteroides thetaiotaomicron VPI-5482* | BT_4562 | BT_4563 | no | BT_4000 | BT_0154 | BT_0576 |
| *Porphyromonas gingivalis W83* | no | no | no | PG_0140 | no | PG_2149 |
| *Prevotella intermedia ATCC 15033* | no | no | no | D2S45_03930 | no | D2S45_01425 |
| *Sphingobacterium spiritivorum ATCC 33861* | no | no | no | HMPREF0766_11792 | no | no |
| *Aureispira sp. EL160426* | no | no | no | AsAng_0024380 | no | no |
| *Gracilimonas mengyeensis DSM 21985* | no | no | no | SAMN06265219_11337; SAMN06265219_10168 | no | no |
| *Ignavibacterium album JCM 16511* | no | no | no | IALB_1374 ; IALB_1587 | no | no |
| *Salinibacter ruber DSM 13855* | no | no | no | SRU_0516; SRU_0157 | no | no |
| *Chlorobaculum tepidum DSM 12025* | no | no | no | no | no | no |
| *Fibrobacter succinogenes S85* | no | no | no | Fisuc_0481; Fisuc_0482; Fisuc_2375 | no | no |
| *Gemmatimonas aurantiaca strain DSM 14586* | no | no | no | GAU_1575; GAU_1564 | no | no |
| *Caldithrix abyssi DSM 13497* | no | no | no | Calab_1398; Calab_1469 | no | no |

**Table S2│Bacteria strains used in this study.**

| Strain | Genotype and Details | Reference |
|---|---|---|
| ***E. coli*** | | |
| NEB® 5-α | *fhuA2 Δ(argF-lacZ)U169 phoA glnV44 Φ80 Δ(lacZ)M15 gyrA96 recA1 relA1 endA1 thi-1 hsdR17* | NEB |
| DH5α λpir | F- *φ80 lac ZΔM15 Δ (lacZYA-arg F)* λ*pir U169 endA1 recA1 hsdR17 (rk-,mk+) supE44λ- thi -1 gyrA96 relA1 phoA* | [1] |
| HB101 | $F^{-}$ Δ*(gpt-proA)62 leuB6 glnV44 ara-14 galK2 lacY1* Δ*(mcrC-mrr) rpsL20* ($Str^{R}$) *xyl-5 mtl-1 recA13 thi-1 hsdSB20* $\lambda^{-}$ | [2] |
| S17-1 λpir | *TpR SmR recA thi pro hsdR-M+RP4:2- Tc:Mu:Km* Tn7 *λpir.* | ATCC |
| BL21 (DE3) | $F^{-}$ *ompT* $hsdS_B$ ($r_B^-$, $m_B^-$) *galdcmrne131 (DE3)* | Invitrogen |
| ***B. thetaiotaomicron*** | | |
| VPI-5482 $tdK^{-}$ | Δ*tdK* | [3] |
| XLBT_001 | VPI-5482 Δ*tdK twinstrep-lptD* | This study |
| ***F. johnsoniae*** | | |
| UW101 | | [4] |
| XLFJ_000 | UW101 Δ*fjoh_1945* | [5] |
| XLFJ_1374 | XLFJ_000 *HA-lptD* | This study |
| XLFJ_1375 | XLFJ_000 *twinstrep-lptD* Δ*lptR* | This study |
| XLFJ_1378 | XLFJ_000 *twinstrep-lptD* Δ*lptQ* | This study |
| XLFJ_1379 | XLFJ_000 *twinstrep-lptD* Δ*lptS* | This study |
| XLFJ_1380 | XLFJ_000 *twinstrep-lptD* Δ*lptT* | This study |
| XLFJ_1391 | XLFJ_000 $P_{ompA}$-*tetR* $P_{ompAinduc}$-*HA-lptD* | This study |
| XLFJ_1404 | XLFJ_000 $P_{ompA}$-*tetR* $P_{ompAinduc}$-*HA-lptD* Δ*lptD* | This study |
| XLFJ_1405 | XLFJ_000 $P_{ompA}$-*tetR* $P_{ompAinduc}$-*HA-lptD* Δ*lptD ALFA-lptN* | This study |
| XLFJ_1409 | XLFJ_000 *HA-lptD* $P_{ompA}$-*tetR* $P_{ompAinduc}$-*ALFA-lptN* | This study |
| XLFJ_1410 | XLFJ_000 *HA-lptD* $P_{ompA}$-*tetR* $P_{ompAinduc}$-*ALFA-lptN* Δ*lptN* | This study |
| XLFJ_1415 | XLFJ_000 *HA-lptD ALFA-lptN* | This study |
| XLFJ_1423 | XLFJ_000 *HA-lptD twinstrep-lptN* | This study |

| | | |
|---|---|---|
| sLO55 | XLFJ_000 *twinstrep SusC* | This study |
| sLO100 | XLFJ_000 *twinstrep SusC ∆lptP* | This study |
| sLO108 | XLFJ_000 *twinstrep-lptD* | This study |
| sLO114 | XLFJ_000 *twinstrep-lptD ∆lptP* | This study |
| sLO126 | XLFJ_000 *twinstrep SusC ∆lptQ* | This study |
| sLO127 | XLFJ_000 *twinstrep SusC ∆lptR* | This study |
| sLO128 | XLFJ_000 *twinstrep SusC ∆lptS* | This study |
| sLO129 | XLFJ_000 *twinstrep SusC ∆lptT* | This study |
| sLO162 | XLFJ_000 *twinstrep SusC ∆lptT2* | This study |
| sLO163 | XLFJ_000 *twinstrep LptD ∆lptT* | This study |

**Table S3│Plasmids used in this study.**

| plasmid | Descriptions | Reference |
|---|---|---|
| pYT313 | *sacB* containing mobilizable suicide vector; $Ap^R$ ($Em^R$) | 6 |
| pCP11 | *E. coli-F. johnsoniae* shuttle plasmid; Apr (Emr) | 7 |
| pExchange | Plasmid for Bacteroides spp. genome engineering by allelic exchange. | 3 |
| pRK2013 | Helper plasmid for triparental conjugation; IncP $Tra^+$ $Km^r$ | 8 |
| pET21a(+) | *E. coli* expression vector, T7 promoter; $Ap^r$ | Merck |
| pXL184 | pCP11-derived plasmid used to express C-terminally his-tagged SusE under the control of the *ompA* promoter | 5 |
| pXL328 | pYT313-derived construct used to generate the deletion of *fjoh_0663* ($lptP_{Fj}$) | This study |
| pXL345 | pYT313-derived construct used to generate the deletion of *fjoh_0204* ($lptT_{Fj}$) | This study |
| pXL346 | pYT313-derived construct used to generate the deletion of *fjoh_2499* ($lptR_{Fj}$) | This study |
| pXL347 | pYT313-derived construct used to generate the deletion of *fjoh_2498* ($lptQ_{Fj}$) | This study |
| pXL348 | pYT313-derived construct used to generate the deletion of *fjoh_2499* ($lptS_{Fj}$) | This study |
| pXL351 | pYT313-derived construct used to generate the N-terminally HA-tagged $lptD_{Fj}$ | This study |
| pXL360 | pYT313-derived construct used to generate the N-terminally ALFA-tagged $LptN_{\mathrm{Fj}}$ | This study |
| pXL367 | pYT313-derived construct used to generate the strain expressing HA-LptD under the control of the aTC-inducible $P_{ompAinduc}$ promoter | This study |
| pXL370 | pYT313-derived construct used to generate the strain expressing ALFA-LptN under the control of the aTC-inducible $P_{ompAinduc}$ promoter | This study |
| pXL383 | pYT313 derived construct used to generate the N-terminally twinstrep-tagged $LptN_{\mathrm{Fj}}$ strain | This study |
| pXL391 | pExchange-derived construct used to generate the N-terminally twinstrep-tagged $lptD_{Bt}$ | This study |
| pXL393 | pCP11-derived plasmid used to express C-terminally FLAG-his-tagged SusG (Fjoh_1210) under the control of the *ompA* promoter | This study |
| pXL409 | pCP11-derived plasmid used to express C-terminally FLAG-his-tagged PorO (Fjoh_4270) under the control of the *remA* promoter | This study |
| pLO123 | pYT313-derived construct used to generate the deletion of $lptD_{Fj}$ | This study |
| pLO128 | pYT313-derived construct used to generate N-terminally twinstrep tagged $lptD_{Fj}$ | This study |

| | | |
|---|---|---|
| pLO137 | pET21-derived vector to heterologously overproduce signal peptide-less Fjoh_0663 (LptP) with a N-terminal 8His tag | This study |
| pLO138 | pET21-derived vector to heterologously overproduce signal peptide-less Fjoh_1847 (LptE) with a N-terminal 8His tag | This study |
| pLO140 | pET21-derived vector to heterologously overproduce signal peptide-less Fjoh_2499 (LptR) with a N-terminal 8His tag | This study |
| pLO144 | pET21-derived vector to heterologously overproduce signal peptide-less Fjoh_2510 (LptS) with a N-terminal 8His tag | This study |
| pLO155 | pYT313-derived construct used to generate the deletion of *fjoh_1207* (*lptT2*) | This study |

**Table S4│Primers used in this study.**

| oligos | Sequences (5′ to 3′) |
|---|---|
| priAK0113 | GATCGTTGTCAGAAGTAAGTTGGCCGCAGTGTTATCACTCATGGT |
| priAK0114 | ACCATGAGTGATAACACTGCGGCCAACTTACTTCTGACAACGATC |
| priXL01 | GGATCCTCTAGAGTCGACCT |
| priXL02 | CCCGAATTTTTCCGCTGCAT |
| priXL05 | CACCAAGGAAAGTCTACACGA |
| priXL06 | CAGGAAACAGCTATGACCAT |
| priXL51 | GTCTACACGAACCCTTTGG |
| priXL52 | TCCACTTTCATATTACGGTTTA |
| prixl177 | AGTAGGCGTGGTATTAAATTT |
| priXL515 | CATCTTAATTTTTTTAATTACAATTTAGTTAATTCTCTATCA |
| priXL536 | TCAATGTTCTGATGGTGTTAACTG |
| priXL721 | CTTGCATGCCTGCAGGTCGACTCTAGAGGATCCCAAATGAGTATAATCAGGCAG |
| priXL722 | CGTCTTTTCTCTTGCAGCTCATCATTAATAATACCAA |
| priXL723 | ATTATTAATGATGAGCTGCAAGAGAAAAGACGACGAAAAG |
| priXL724 | CCGAAGCAGGGTTATGCAGCGGAAAAATTCGGGGTTTTTTATTGGCGAATCAAA |
| priXL781 | AAGCTTGCATGCCTGCAGGTCGACTCTAGAGGATCCAAAGGTGCCGTAAACTATATTA |
| priXL782 | GTTACATCGTTGGGATAAATGACAATTTTATTCACTATCCGTT |
| priXL783 | CGGATAGTGAATAAAATTGTCATTTATCCCAACGATGTAACAT |
| priXL784 | ACCCCGAAGCAGGGTTATGCAGCGGAAAAATTCGGGAGCAGTAAAGTCCACCAAGTT |
| priXL787 | AAGCTTGCATGCCTGCAGGTCGACTCTAGAGGATCCCAATTGCTAAGGCTTCTAAAAA |
| priXL788 | CAGGCGCAGAGTAATAGAATATTCTCATTTGATAGCGTTA |
| priXL789 | CTATCAAATGAGAATATTCTATTACTCTGCGCCTGATAATC |
| priXL790 | ACCCCGAAGCAGGGTTATGCAGCGGAAAAATTCGGGGCATCCAGTCATCATTAAACA |
| priXL793 | AAGCTTGCATGCCTGCAGGTCGACTCTAGAGGATCCTTTGTCCCTTTGTCACTTTGCA |
| priXL794 | GTATACTTTTAGTCTTATCATAAACCAAGCTCCAAAAA |
| priXL795 | TTGGAGCTTGGTTTATGATAAGACTAAAAGTATACTTTAACGTG |
| priXL796 | ACCCCGAAGCAGGGTTATGCAGCGGAAAAATTCGGGATTGCCGGAATTTGAAACTAT |
| priXL799 | TTATAAAACTGTAAATGAAAGATAATACAGCCAATTCAGGA |

| | |
|---|---|
| priXL800 | GAATTGGCTGTATTATCTTTCATTTACAGTTTTATAATTATCGAAGT |
| priXL801 | AAGCTTGCATGCCTGCAGGTCGACTCTAGAGGATCCCCGGATACATATAAAATGCTG |
| priXL802 | TGCGACGATGGTACAGTAAACTTATAGTAAAAACAAAAGGATTC |
| priXL803 | TTTTTACTATAAGTTTACTGTACCATCGTCGCAGCCGTTTAA |
| priXL804 | ACCCCGAAGCAGGGTTATGCAGCGGAAAAATTCGGGCTGAAGAAAATGGCAGTGTAA |
| priXL813 | AGCGTAATCTGGAACATCGTATGGGTAAGCAGGTAAAGATTTTTTTTTAGGTTT |
| priXL814 | CTGCTTACCCATACGATGTTCCAGATTACGCTGTAAAACAAACAAATAAGCCGGC |
| priXL815 | GAATTAACTAAATTGTAATTAAAAAAATTAAGATGATTATTTTTGCCAAAAAATTATAT |
| priXL816 | TCTTCAGTGCAGTTAACACCATCAGAACATTGATTATCGATTTATAGTGCTTCGT |
| priXL831 | CATCACCATCATCACCACCA |
| priXL832 | ACTTAATTTTTTTAATTACAATTTAGTTAATTAC |
| priXL840 | ATATACAGGTCCTTATGGAGT |
| priXL841 | AAACTTATAATCTGAGGCTGG |
| priXL842 | AAATGATTGAAGATGCTGAGG |
| priXL843 | CAATTATAGTTCCTCCTGGTT |
| priXL844 | GTTAATCTACGTCTTAATTCCTCTTCTAATCTAGAACCTTTTGAAGTAGTTTTTTCTTG |
| priXL845 | TCTAGATTAGAAGAGGAATTAAGACGTAGATTAACAGAAAGTTTAGATATGAATACAAAATATTCAA<br>G |
| priXL854 | GAATTAACTAAATTGTAATTAAAAAAATTAAGATGAGAATATTCATTTTTTTATATCTATTAATTATAC |
| priXL855 | CTTCTTCAGTGCAGTTAACACCATCAGAACATTGATTATTGGAAGAAATTCCAAACTAA |
| priXL860 | GAGAATTAACTAAATTGTAATTAAAAAAATTAAGATGCATACAAACTTATTTAATATCG |
| priXL872 | ACTCTTCGGGATGCAAAAATA |
| priXL873 | AGAGAGAAACATCTTTTGCATAC |
| priXL876 | CGGTCGGAAAAGGTATGCAAAAGATGTTTCTCTCTGAACTTCTGGCTTGCATATCA |
| priXL879 | TGGAGTTATTTTCATATTTTTGCATCCCGAAGAGTTGCAATGTACCGGAGTATTTG |
| priXL912 | CGCCGAACCTCCCGATCCACCTCCGGAACCTCCACCTTTTTCGAACTGCGGGTGGCTCCAACC<br>TTTTGAAGTAGTTTTTTCTTG |
| priXL913 | GGTGGAGGTTCCGGAGGTGGATCGGGAGGTTCGGCGTGGTCACATCCACAATTTGAGAAGAG<br>TTTAGATATGAATACAAAATAT |
| priXL922 | TAATTAACTAAATTGTAATTAAAAAAATTAAGTATGAAATCAAATATAAAGATCGTTT |

| | |
|---|---|
| priXL923 | GAATTTTAGTGATGGTGGTGATGATGGTGATGATTTTTCAATATTATGTATTGAAAAGG |
| priXL926 | AATTATGCAGTGCTGCCATAA |
| priXL927 | CTCCGATCGTTGTCAGAAGTAA |
| priXL930 | GGTGGAGGTTCCGGAGGTGGATCGGGAGGTTCGGCGTGGTCACATCCACAATTTGAGAAGGT TGACTCTGTAGCACCGAC |
| priXL940 | GAATTTTAGTGATGGTGGTGATGATGGTGATGTTTGTCATCGTCATCTTTATAATCATTTTTCAATA TTATGTATTGAAAAGG |
| priXL960 | CAAGCTTGCATGCCTGCAGGTCGACTCTAGAGGATCCGGACAATCTTTTTGGATGAAG |
| priXL961 | CCAAAAAATTACCATTTTGCCAGTGACTCATTAAAAAAAGTGGTTGCGACTAATAATAG |
| priXL962 | AAATATCTTTTACTATTATTAGTCGCAACCACTTTTTTTAATGAGTCACTGGCAAAA |
| priXL963 | GACCCCGAAGCAGGGTTATGCAGCGGAAAAATTCGGGTTTCTTCTTCACTTAGTGCAG |
| priXL998 | GATGACGATGACAAACATCACCATCATCACCACCATCACTAAAATTCAGCAATAGAAACCTCCAA |
| priXL999 | CACCCAAAAACAAAAATTTAATACCACGCCTACTATGCGTCAATACTTTACAATC |
| priXL1000 | TGGTGATGATGGTGATGTTTGTCATCGTCATCTTTATAATCTTTAGAAGGGAAGTCTTTACT |
| oLO334 | GCCTCTCCCCGCGCGGTTAAGTGTTACTGGTGCTGAAA |
| oLO335 | CCATAGGCTGCCAGTTTAAATAAGTTTGTATGCAAAGGTTTAAATACTA |
| oLO336 | TACAAACTTATTTAAACTGGCAGCCTATGGG |
| oLO337 | TTGGATACACCAAGGCTTAAAACAAATCTTTCAGGAGTATTTTACT |
| oLO338 | CACGAGGGAGCTTCCAGG |
| oLO350 | GCCTCTCCCCGCGCGCCCTTTAAAAAGCTGTATCCCC |
| oLO351 | GTGGCTCCAAGCGCTAGCAGGTAAAGATTTTTTTTTAGGTTT |
| oLO352 | AAATCTTTACCTGCTAGCGCTTGGAGCC |
| oLO353 | TTGGATACACCAAGGATCAAAACCATTTACGGTTGTG |
| oLO364 | ATTTGTTTGTTTTACACTAGTTTTCTCGAACTGCG |
| oLO365 | TTCGAGAAAACTAGTGTAAAACAAACAAATAAGCCGGC |
| oLO397 | TCTGGCGGTGGATCGAGCATTACCGGCGGA |
| oLO398 | AGCCGGATCTCATCATTAGAAAGCTTTTCGTTTCTTTTCG |
| oLO399 | TCTGGCGGTGGATCGGGCGTTTATAACTTTACAGGAGC |
| oLO400 | AGCCGGATCTCATCATTACCATTTTGCCAGTGACTC |
| oLO407 | TCTGGCGGTGGATCGGCAAATACGCCTACAACTCC |

| | |
|---|---|
| oLO408 | AGCCGGATCTCATCACTAAAAGTATACTTTAACGTGTGTTTCA |
| oLO414 | TTCTGGCGGTGGATCGGACGATGGTGATCTAACCG |
| oLO415 | GCCGGATCTCATCACTATAAGTTTACTGTTATTTTACCCAAAG |
| oLO445 | TCGCTGCGAAAAAACATAAAGCTGGCTTTGAAAATTTTG |
| oLO446 | CAAAGCCAGCTTTATGTTTTTTCGCAGCGAATCG |
| oLO448 | AAATCCCCAAAATAGAAAGCG |
| oLO449 | ATCTCATTTAGTTTAAAAATTAAAAGGCG |
| oLO456 | TCCTCTTAAACCCGGACTAAC |
| oLO457 | GCGGATACAACGGATTTTTATTC |
| oLO458 | TCGAGATCTTTACCTGCTAGTTC |
| oLO459 | CAGCAGGAATTGTTGGACC |
| oLO475 | AGGTCGACTCTAGAGGATCCAATCTTGTTTTGAATATGATAAAGATGGAG |
| oLO476 | ATGCAGCGGAAAAATTCGGGATTTCACCTGCAAATGCAAC |
| oLO527 | AAAAGAAGGAATAACACAAACCTG |
| oLO528 | GATAAAAATGAAATGTTGGAAACGAG |
| oLO529 | TCCTGTTCTACTTTTGTCAAACTTG |
| oLO530 | ATTGGCCAAAAGTGCGTAC |
